# Taxonomic and carbon metabolic diversification of Bathyarchaeia during its co-evolution history with the early Earth surface environment

**DOI:** 10.1101/2022.11.19.517056

**Authors:** Jialin Hou, Yinzhao Wang, Pengfei Zhu, Na Yang, Lewen Liang, Tiantian Yu, Mingyang Niu, Kurt Konhauser, Fengping Wang

## Abstract

Bathyarchaeia, as one of the most abundant microorganisms on Earth, play vital roles in the global carbon cycle. However, our understanding of their origin, evolution and ecological functions remains poorly constrained. Based on the phylogeny of the present largest dataset of Bathyarchaeia metagenome assembled genome (MAG), we reclassified Bathyarchaeia into eight order-level units and corresponded to the former subgroup system. Highly diversified and versatile carbon metabolisms were discovered among different orders, particularly atypical C1 metabolic pathways, indicating that Bathyarchaeia represent overlooked important methylotrophs. Molecular dating results indicate that Bathyarchaeia diverged at ∼3.3 Ga, followed by three major diversifications at ∼3.0 Ga, ∼2.5 Ga and ∼1.8-1.7 Ga, likely driven by continental emergence, growth and intensive submarine volcanism, respectively. The lignin-degrading Bathyarchaeia clade emerged at ∼300 Ma and perhaps contributed to the sharply decreased carbon sequestration rate during the Late Carboniferous period. The evolutionary pathway of Bathyarchaeia potentially have been shaped by geological forces, which in turn impacted the Earth’s surface environment.

**Teaser:** The origin and divergence of Bathyarchaeia linked to the early Earth tectonics and surface environment changes

## Introduction

Bathyarchaeia, previously known as the Miscellaneous Crenarchaeotal Group (MCG) (*1*) or Bathyarchaeota (*2*), are ubiquitously distributed dominating archaeal communities in anoxic subsurface environments, including oceanic (*3*), freshwater (*4*), hydrothermal (*5*), hot spring sediment (*6*) and soil (*7*), in particular coastal settings (*8, 9*). Globally, it is estimated that Bathyarchaeia account for ∼2.0-3.9 × 10^28^ cells (*5*), ranking as one of the most abundant microbial groups on Earth. However, because there is still no laboratory cultivation of Bathyarchaeia thus far, their taxonomy was defined by the identity of 16S rRNA genes retrieved directly from environmental samples and metagenome-assembled genomes (MAGs). This has resulted in notable phylogenetic inconsistencies and instability. For example, Bathyarchaeia were initially classified into 17 subgroups (*8*) and further extended to 25 subgroups with accumulating sequences (*4, 10*). Recently, the Genome Taxonomy Database (GTDB) archaeal taxonomy reclassified the phylum Bathyarchaeota to a class-level clade and renamed it the class Bathyarchaeia within the phylum Thermoproteota, corresponding to the previous Thaumarchaeota-Aigarchaeota-Crenarchaeota-Korarchaeota (TACK) superphylum (*11*). However, the correspondence between the former subgroup systems and the detailed taxonomy remains unknown, thus impeding our understanding of the evolution and ecological roles of different Bathyarchaeia lineages.

Bathyarchaeia have been suggested having diversified capabilities in metabolizing a large spectrum of organic compounds to fulfill their carbon or energy demands, including peptides/proteins (*3*), polymeric carbohydrates (*12*), fatty acids (*13*), lignin (*14*), aromatic (*2*) and aldehyde compounds (*15*). At the same time, isotopic and enzymatic experiments have demonstrated their ability to assimilate inorganic carbon (*5, 14*). Furthermore, recent findings imply that some Bathyarchaeia species might also undergo anaerobic methane or multiple-carbon alkane metabolism (*16, 17*). Overall, given their numerical dominance and metabolic versatility in the anoxic deep biosphere, different Bathyarchaeia lineages are believed to be critical players in modern carbon cycling. Nevertheless, when and how Bathyarchaeia originated on early Earth and subsequently evolved into such a diversified and currently flourishing branch of life remains unclear.

Recently, the evolution of several archaea with special ecological functions, such as ammonia-oxidizing Thaumarchaeota (*18, 19*), methanogens (*20*) and methanotrophic archaea (*21*), has been elucidated by molecular dating analysis. To demonstrate the metabolic characteristics and evolutionary history of Bathyarchaeia, we propose here an improved taxonomy derived from an exhaustive collection of 304 non-redundant representative MAGs refined from available public databases and our own datasets based on the GTDB criterion (*11*). This approach enabled a comprehensive investigation of the distinctive environmental distribution and taxonomy-specific carbon metabolic traits of different Bathyarchaeia lineages. Their origin and divergence time were inferred from robust phylogenomic and molecular dating analysis, reflecting the multiple independent correlations between the evolution of Bathyarchaeia and key geological processes that occurred in the early Earth’s history, including subaerial continental production, growth and episodes of intensive submarine volcanism. In addition, we propose that the origin of lignin-degrading Bathyarchaeia clade probably contributed to the contemporaneous sharp decrease in coal deposition rate during the Late Carboniferous period.

## Results and discussion

### Improved Bathyarchaeia taxonomy and subgroup assignment

A total of 304 non-redundant representative Bathyarchaeia MAGs identified from available public genome databases, including the National Center for Biotechnology Information (NCBI), the genomic catalog of Earth’s microbiome (GEM) (*22*) and GTDB (*11*), as well as our sequenced datasets (Table S1), were utilized for further phylogenomic analysis (Table S2). A total of 78 (25.7%) and 187 (61.5%) MAGs had high (completeness ≥ 90% and contamination < 5%) and medium (completeness ≥ 50% and contamination < 10%) genomic quality, respectively (Table S3). Based on the robust phylogenomic inference, we propose a systematic taxonomy for each Bathyarchaeia representative MAG, whose taxonomic rank was hierarchically assigned and normalized by the approach implemented in GTDB (*23*). First, the calculated relative evolutionary divergence (RED) value (0.339) for the whole Bathyarchaeia lineage indicates it as an independent class-level taxonomic unit within the phylum Thermoproteota (previously known as TACK superphylum), in accordance with the archaeal GTDB taxonomy (Release 06-RS202) (Fig. S1 and Table S4). Furthermore, the 304 representative MAGs were classified into eight order-level lineages and corresponding subsequent family and genus ranks (Table S3 and S4).

The eight Bathyarchaeia orders were named after different Chinese traditional mythological figures, denoting their distinct environmental features. These include Bifangarchaeales (Bifang, a single-legged bird as the symbol of wildfire, this lineage largely comes from hot springs); Jinwuousiales (Jinwu, a three-legged bird living in the Sun, their members are mostly from hydrothermal vents); Zhuquarculales (Zhuque, a red bird as the god of fire, all the species are from hydrothermal vents); Wuzhiqibiales (Wuzhiqi, the god of Huai River, the majority comes from marine sediments, in particular estuaries); Houtuarculales (Houtu, the god of earth, large parts are discovered from soils); Mazuousiales (the sea goddess, mostly comes from East China Sea sediments); Baizomonadales (Baize, a propitious creature living in cold Kunlun Mountains, this lineage has miscellaneous distribution but largely from cold marine sediments); and Xuanwuarculales (Xuanwu, a black turtle as the god of iciness and water, only has one MAG from underground water sediment) (Fig. 1, see Nomenclature and Etymology and Table S5 for details). Despite using the same RED intervals for taxonomic rank classification, compared with the reference GTDB taxonomy (Release 06-RS202), the newly improved taxonomy of Bathyarchaeia has more designated families and genera for the majority of orders (Table 1). Notably, the order Mazuousiales is a novel Bathyarchaeia order proposed in this study, in which five of their seven MAGs are additionally retrieved from the GEM and our datasets (Table S3). Consequently, these taxonomic differences can be attributed to the largely increased Bathyarchaeia MAGs analyzed here (304 MAGs) in comparison to the reference GTDB database (173 MAGs, Release 06-RS202) (Table S2).

**Fig. 1.**
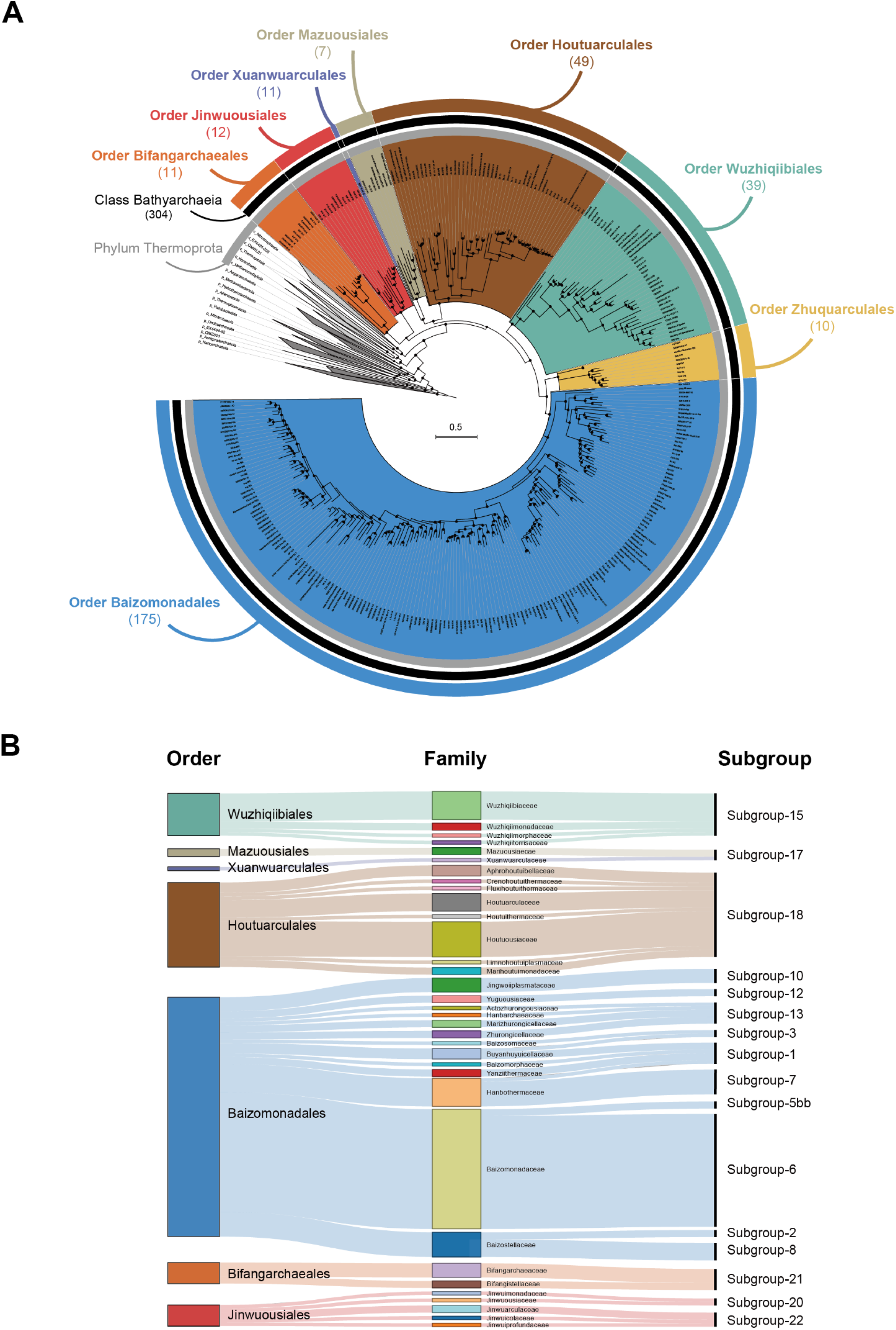
The newly improved taxonomy and subgroup assignment of the class Bathyarchaeia. **(A)**, Phylogenetic tree of 304 representative Bathyarchaeia MAGs refined from publicly available databases and our lab datasets based on a concatenated alignment of 122-archaeal marker proteins implemented in GTDB-tk by using IQ-TREE2 with best-fit *LG+F+R10+C60* model and SH approximate likelihood ratio test with 1000 bootstrap replicates (bootstrap higher than 0.9 are shown with black dots). The colored background and outer rings denote the eight proposed orders of class Bathyarchaeia, and the inner black and gray rings represent the whole class Bathyarchaeia and phylum Thermoproteota, respectively. The number of MAGs used in the phylogenetic analysis is listed under the name for each taxonomic lineage. **(B)**, Taxonomic assignment of subgroups for the representative Bathyarchaeia MAGs with 16S rRNA genes. The phylogenetic tree was constructed with subgroup-classified sequences from Zhou et al. (*10*) as a reference by using raxML 8.2.12 with *-m GTRGAMMA -N autoMRE* and 1000 bootstrap replicates (Fig. S1).

**Table 1.**
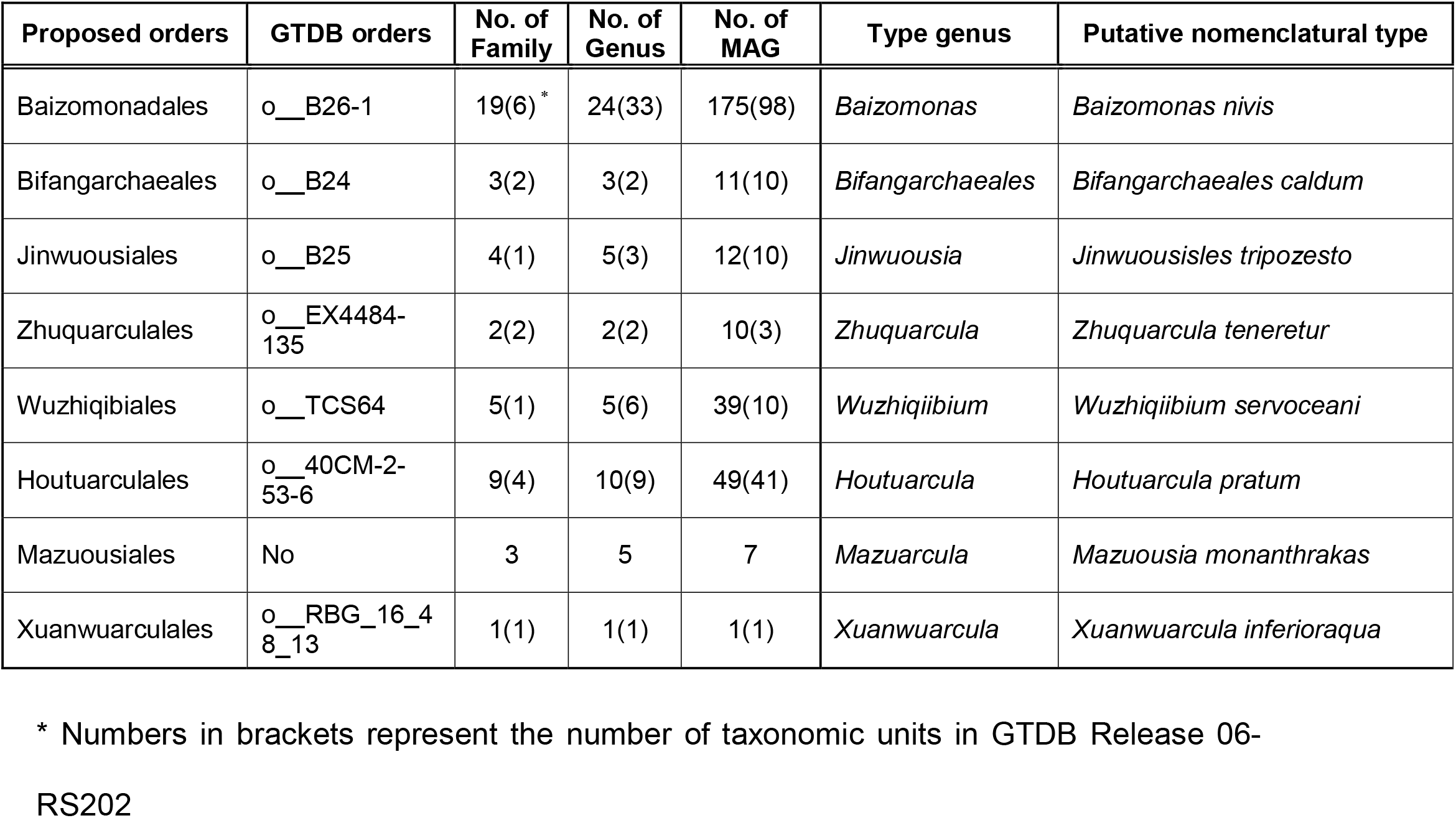
The newly improved taxonomy of the class Bathyarchaeia in this study and the difference from the reference GTDB taxonomy (Release 06-RS202)

Previously, Bathyarchaeia were classified into 25 subgroups based on 16S rRNA gene phylogeny. However, subgroup is not a legitimate rank name and normally subjectively defined without a consistent phylogenetic threshold (*4, 8, 10*), e.g., the minimum intragroup identities of different subgroups range from 84%-94% in the initial designation (*8*) and subsequently were updated to >90% (*10*), which generally fall into the universal taxonomic threshold range between the order and family level (89.2%-92.25% (*24*)). As a consequence, the specific taxonomic rank for each subgroup and whether they are equivalent remain unresolved. To address this, a phylogenetic tree was constructed using the 16S rRNA genes retrieved from the representative Bathyarchaeia MAGs and subgroup-classified reference sequences (*10*) to establish the links between each Bathyarchaeia subgroup and their corresponding taxonomic ranks (Fig. S2). Except for the order Zhuquarculales (as no 16S rRNA gene was retrieved from their MAGs), the other seven Bathyarchaeia orders have at least one MAG encoding the 16S rRNA gene with subgroup assignment (Fig. 1b).

Overall, the 16S rRNA genes from 16 of the 25 Bathyarchaeia subgroups were identified in the representative Bathyarchaeia MAGs (Fig. 1b). Specifically, subgroups-15, -18 and -21 correspond to the orders Wuzhiqibiales, Houtuarculales and Bifangarchaeales, respectively; subgroup-17 is assigned to the orders Xuanwuarculales and Mazuousiales; and the order Jinwuousiales contains the MAGs belonging to subgroup-20 and -22. For the largest order, Baizomonadales, 175 MAGs were distributed into ten previous subgroups, including subgroup-1, -2, -3, -5, -6, -7, -8, -10, -12 and -13 (Fig. 1b), suggesting that these subgroups actually correspond to taxonomic ranks lower than the order level. Intriguingly, subgroup-8 is one of the most dominant Bathyarchaeia lineages in marine sediments (*25, 26*). Here, all five MAGs with subgroup-8 16S rRNA genes were monophyletically assigned to the genus *Baizosediminiarchaeum* within the order Baizomonadales, indicating that this important Bathyarchaeia subgroup very likely belongs to a genus-level taxonomic unit. In addition, as the largest genus of the class Bathyarchaeia, *Baizomonas* contains the MAGs assigned to subgroup-6 and -5bb, which are frequently detected in freshwater sediments (*4*). In summary, these results demonstrate that the different Bathyarchaeia subgroups assigned previously were far from taxonomic equivalents. Instead, they correspond to remarkably distinct ranks in the improved Bathyarchaeia taxonomy, ranging from the above order level to the below genus level. As a consequence, cautions should be applied when extrapolating the significance of different Bathyarchaeia subgroups, as they may not be suitable for quantitative comparisons either among themselves or with other microbial lineages.

### Taxonomically specific distribution in terrestrial and geothermal environments

Nearly 50% of the Bathyarchaeia MAGs (150) were isolated from oceanic environments, of which 91% were recovered from marine sediments that were associated with the seafloor (70) or submarine hydrothermal systems (67), while the rest were found in seawater (8) and crustal fluids (5) (Fig. 2a). The other 50% of the MAGs are from terrestrial ecosystems, including soils (66), hot springs (66) and freshwater sediments (17). In particular, eight of eleven MAGs from the order Bifangarchaeales were recovered from terrestrial hot spring sediments, while the order Houtuarculales contained ∼61% (30 in 49) MAGs from soils, especially the genus *Houtuousia* (all of 23) (Fig. 2b and Table S3). Therefore, these Bathyarchaeia lineages may already have specifically adapted to different terrestrial ecosystems, which expand the range of ecological impacts and significance of the class Bathyarchaeia from oceanic to terrestrial environments. Besides, only one MAG from the order Xuanwuarculales was recovered from the underground water sediment (*27*) (Fig. 2b), implying that more sampling efforts are needed to explore the potential novel Bathyarchaeia species inhabiting the deep terrestrial biosphere. Taken together, over 75% of Bathyarchaeia MAGs (228) were retrieved from diverse sedimentary environments in both oceanic and terrestrial ecosystems.

**Fig. 2.**
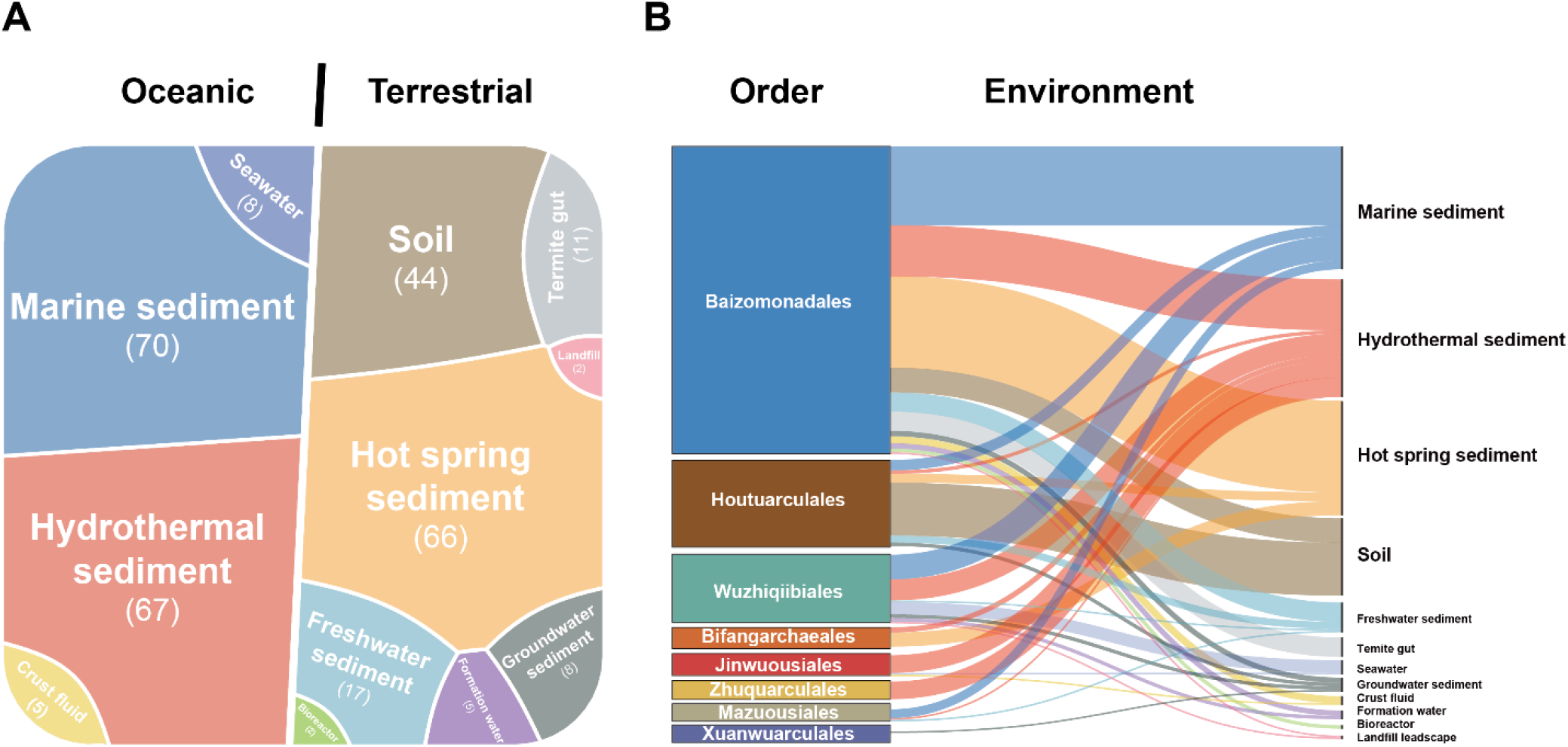
Environmental distribution of the class Bathyarchaeia. **(A)**, Distribution of 304 representative Bathyarchaeia MAGs across different ecological types from oceanic and terrestrial environments. The number of MAGs associated with each ecological type is indicated below the name. **(B)**, Environmental distribution of representative Bathyarchaeia MAGs from each proposed order.

Except for the predominant Bathyarchaeia groups in the temperate coastal zone or cold deep subsurface (*3, 8, 10*), diverse Bathyarchaeia species have also been discovered in different geothermal environments (*5, 6*). Here, a total of 143 (∼47%) Bathyarchaeia MAGs were retrieved from hot springs (66), hydrothermal sediments (67), crustal fluids (5) and deep formation waters (5) (Fig. 2a), along with significant taxonomic specificity at the order level. For instance, all MAGs of the orders Bifangarchaeales, Zhuquarculales and Jinwuousiales (except one from saline water) were specifically recovered from geothermal environments (Fig. 2b and Table S3), consistent with previous subgroup-based studies (*5, 6, 10, 28*). In particular, eight of eleven MAGs from the order Bifangarchaeales were exclusively recovered from hot springs, whereas the other two orders only contain the MAGs from hydrothermal sediments (Fig. 2b). Collectively, these results indicate that diversified Bathyarchaeia lineages thrive in high-temperature environments and play important roles in global geothermal ecosystems. In fact, all members from the orders Bifangarchaeales, Jinwuousiales and Zhuquarculales are very likely obligate thermophilic archaea that evolved in high-temperature environments, in accordance with previous studies about the thermal origin and adaptation of Bathyarchaeia (*6, 28*).

### Diversified metabolic potentials for complex organic carbon utilization

Metabolic reconstruction of 86 high-quality Bathyarchaeia MAGs indicated that almost all eight proposed orders possess relatively similar and complete central carbon metabolisms, including glycolysis, gluconeogenesis, reverse ribulose monophosphate (RuMP), phosphoribosyl pyrophosphate (PPRP) biosynthesis and pyruvate-ferredoxin oxidoreductase (PFO complex), in addition to complete gene sets coding for the H_4_-MPT Wood-Ljungdahl pathway (WLP) (Fig. 3a and Table S6). In addition, highly diversified carbohydrate-active enzymes (CAZymes) and peptidases were extensively predicted in their MAGs (Fig. 3b and 3c). All these core metabolic repertoires reflect the versatile capabilities of Bathyarchaeia in utilizing different organic compounds. Consistent with this inference, previous studies have demonstrated that different Bathyarchaeia species can assimilate a variety of organic carbon compounds, such as acetate (*13*), amino acids (*29*), fatty acids and complex hydrocarbons (*12*). Some Bathyarchaeia members are also proposed to be capable of utilizing lignin as an energy source for acetogenesis through the reductive acetyl-CoA pathway (also reverse H_4_-MPT WLP) (*5, 14*).

**Fig. 3.**
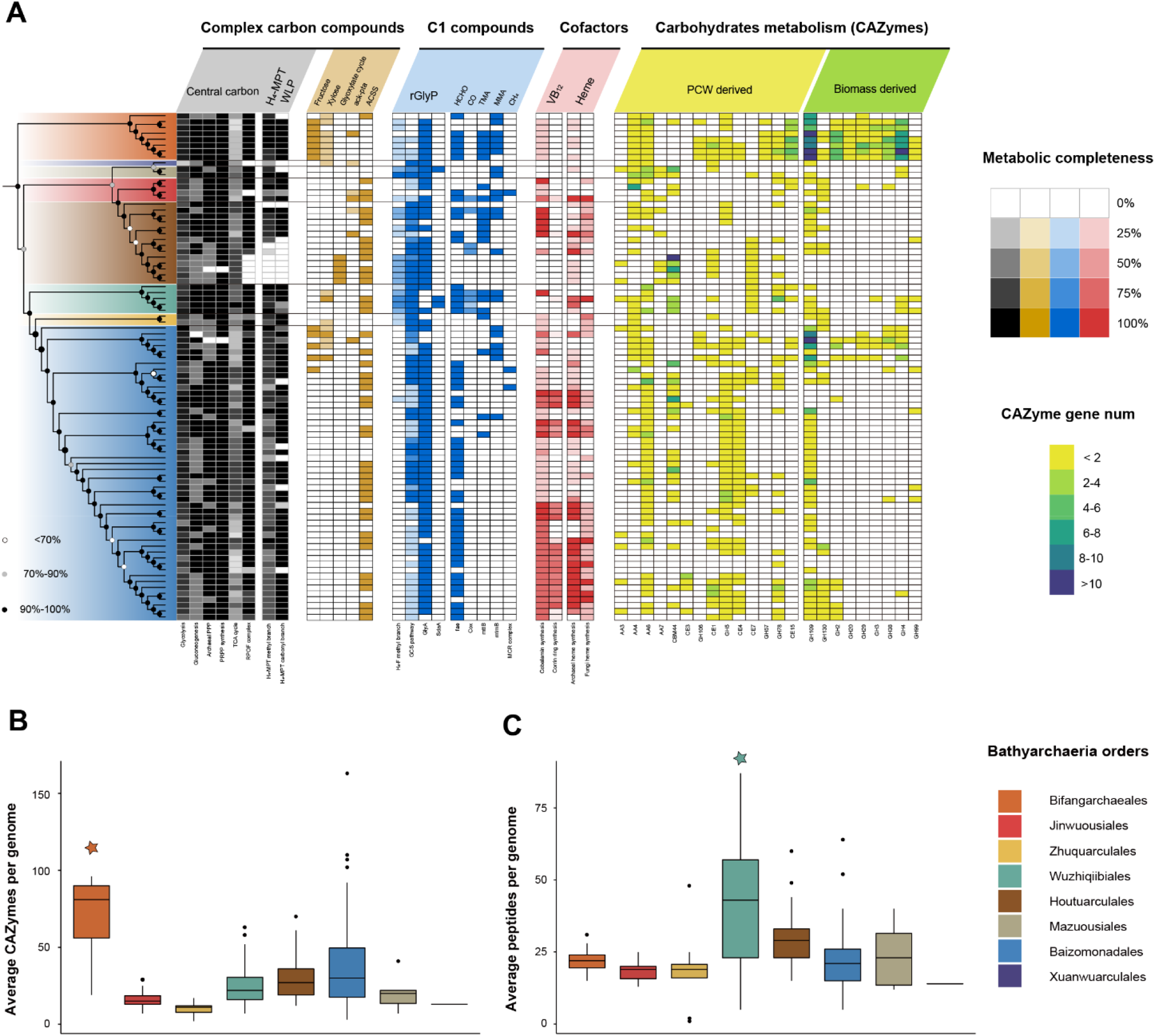
Distinctive metabolic traits of the class Bathyarchaeia. **(A)**, The phylogenomic tree includes 86 high-quality representative Bathyarchaeia MAGs and was inferred by iQ-TREE2 with the best-fit *LG+F+R7+C20* model from 122 archaeal marker proteins implemented in GTDB-Tk (see Method and Materials). Colors on branches indicate eight proposed Bathyarchaeia orders. The completeness of each metabolic pathway is defined by the ratio of marker genes identified in the complete gene repertoire for each MAG, while for many key genes or complexes, such as those involved in C1 compound metabolism, their presence and absence in each MAG are directly indicated by the percentage of completeness. Metabolic capabilities of carbohydrate degradation are evaluated by the gene numbers of different CAZymes identified in each MAG. The following metabolic pathways and marker genes were used in the figure (see Table S6): archaeal PPP, archaeal pentose phosphate pathway; PRPP synthesis, phosphoribosyl pyrophosphate synthesis; TCA cycle, tricarboxylic acid cycle; PFOR complex, pyruvate:ferredoxin oxidoreductase complex; ack-pta, acetate kinase and phosphate acetyltransferase; ACSS, acetyl-CoA synthetase; rGlyP, reductive glycine pathway; GCS, glycine cleavage system; *GlyA*, glycine hydroxymethyltransferase; *sdaA*, L-serine dehydratase; fae, formaldehyde-activating enzyme; *cox*, aerobic carbon-monoxide dehydrogenase; *mttB*, trimethylamine-corrinoid protein Co-methyltransferase; *mtmB*, methylamine-corrinoid protein Co-methyltransferase; MCR complex, methyl-coenzyme M reductase complex; AA, auxiliary activity; GH, glycoside hydrolase; CE, carbohydrate esterase. **(B)** and **(C)**, Box plots show the average number of CAZyme and peptide (MERPO) genes per genome for each Bathyarchaeia order, respectively. The stars indicate that the order has a significantly higher average genomic gene number in comparison with all other Bathyarchaeia orders (Wilcoxon test with fdr correction P < 0.05).

In addition to the common central carbon metabolism, different Bathyarchaeia orders show distinct metabolic preferences in metabolizing complex organic compounds. Among all Bathyarchaeia orders, the hot spring order Bifangarchaeales has the highest average number of CAZyme genes per genome in their MAGs (Fig. 3b), particularly those associated with decomposing biomass (e.g., GH4 (*30*) and GH109 (*31*)) and plant cell wall (PCW)-derived polysaccharides (e.g., AA4 (*32*), AA6 (*33*) and CE15 (*34*)) (Fig. 3a). Moreover, the key genes for fructose and xylose degradation were also predicted in seven of their eight MAGs but rarely detected in others (Fig. 3a). Therefore, these results indicate that the members of the order Bifangarchaeales likely had the capacity to use a large spectrum of carbohydrates, especially those derived from vascular plants, probably for specifically adapting to the hot spring environment, where is usually enriched with vegetal organic deposits (*35*). More importantly, we demonstrate that this ability was specific to the order Bifangarchaeales rather than a common metabolic trait of all thermophilic Bathyarchaeia species at high temperatures, as proposed before (*6*).

For the orders Jinwuousiales and Zhuquarculales, these two hydrothermal lineages have significantly smaller average genome sizes, lower coding densities and fewer gene numbers than the other orders, including the hot spring order Bifangarchaeales (Fig. S3). It means that remarkable genome streamlining might have occurred in these hydrothermal Bathyarchaeia lineages, possibly as a consequence of the extreme conditions associated with hydrothermal activity and scarce external organic inputs in deep-sea environments. In terms of metabolism, four Jinwuousiales MAGs encode the phosphate acetyltransferase (Pta), acetate kinase (Ack) and/or acetyl-CoA synthetase (ACSS) coding genes for acetogenesis/acetate utilization (Fig. 3a), implying that acetate is an important intermediate substrate for Bathyarchaeia species inhabiting hydrothermal sediments. Compared with the order Jinwuousiales, members of the order Zhuquarculales have higher GC content and lower N-ARCS (nitrogen per amino-acid residue side chain) (*36*) (Fig. S3) and few additional genes for amino acid synthesis and CAZymes (Fig. 3a and Fig. S4). These atypical inabilities in terms of both anabolism and catabolism indicate that the order Zhuquarculales might physiologically depend on other organisms for acquiring bioavailable nitrogen sources and other essential nutrients to survive in extreme hydrothermal environments.

Over 72% of the MAGs in the order Houtuarculales are from non-geothermal terrestrial environments (Fig. 2b). Intriguingly, all five MAGs of the genus *Houtuousia*, which were exclusively recovered from grassland soils (Table S3), uniquely encode the two key genes in the glyoxylate cycle, isocitrate lyase and malate synthase, in addition to the relatively complete TCA cycle and ACSS-coding gene (Fig. 3a), which allows them to directly assimilate acetate as a potential carbon source (*37*). Although the glyoxylate cycle has been identified in many bacterial and fungal species, only a few Haloarchaea have been demonstrated to operate this pathway to assimilate acetate (*38*), which might facilitate their adaptation to high-salinity environments (*39*). Here, the glyoxylate cycle discovered in these soil Bathyarchaeia MAGs suggests that the unusual acetate assimilation pathway is not restricted to halophilic archaea, thus expanding its ecological significance from the extreme hypersaline environment to the broader soil environment. Besides, another interesting finding for the genus *Houtuousia* is the absence of the H_4_-MPT WLP and PFOR complex in their MAGs, although these two highly oxygen-sensitive pathways were found in all other Bathyarchaeia MAGs as core metabolic traits (Fig. 3a). In addition, their MAGs also exclusively encode the cytochrome C oxidases putatively for aerobic respiration (Fig. 3a), resembling those in aerobic *Sulfolobus acidocaldarius* (*40*). Therefore, the members of the genus *Houtuousia* probably made physiological adaptations to the (micro)oxic soil environment with a (micro)aerobic lifestyle and atypical acetate assimilation capability.

The order Wuzhiqiibiales solely corresponds to the previous subgroup-15 (Fig. 1b), which frequently predominates the archaeal communities in anoxic coastal sediments (*26*) with positive correlations with salinity and sedimentary depth (*41*). Here, among all eight Bathyarchaeia orders, the MAGs of Wuzhiqiibiales on average encode the highest proportion of genes for peptidases (on average 41.6 genes per genome) (Fig. 3c), especially intracellular peptidases (Fig. S5 and Table S7), indicating their remarkable potential in metabolizing amino acids, peptides and detrital proteins. In accordance with our findings, the first single-cell sequenced Bathyarchaeia genome (SCGC AB-539-E09), assigned to the order Wuzhiqiibiales herein, was shown to play active roles in degrading detrital proteins in anoxic sediments (*3*). Recently, ^13^C-labeled incubation also indicated that some Wuzhiqiibiales species could use extracellular proteins through catabolic processes (*29*). Therefore, using detrital proteins seems to be a metabolic characteristic that is specific to all members of the order Wuzhiqiibiales, reflecting their important roles in protein remineralization and carbon/nitrogen cycling in global marine sediments.

As the most numerous and diversified Bathyarchaeia lineage, 50 MAGs from the order Baizomonadales encode the majority of carbon metabolic pathways identified in other orders (Fig. 3a), and they were retrieved not only from common natural depositional systems but also from a few unique artificial environments, such as bioreactors and landfills (Fig. 2b). This suggests that the order Baizomonadales is currently the most representative and successfully evolved Bathyarchaeia lineage. Intriguingly, the anaerobic cobalamin (vitamin B_12_) synthesis pathway was specifically identified in most members of the genera *Baizomonas* and *Baizosediminiarchaeum* (Fig. 3a and Fig. S4). Despite cobalamin being an essential cobalt-containing cofactor for all three domains of life, in the domain Archaea, only a few Thaumarchaeota (*42*) and euryarchaeotal methanogens (*43*) can *de novo* synthesize it, which might facilitate their key roles in seawater and the methane zone in deep sediments (*44*), respectively. Accordingly, the presence of the cobalamin synthesis pathway in these two genera makes Bathyarchaeia the third archaeal lineage with this unusual capability, suggesting they could serve as a keystone species in the subseafloor microbial community by providing this essential cofactor for other members. In addition, some *Baizosediminiarchaeum* species could grow with lignin as the energy source (*14*) using a novel O-demethylation methyltransferase system that require cobalamin-binding corrinoid protein (*45*), while the growth of certain *Baizomonas* species (assigned to subgroup-6) was also significantly simulated by different lignin-derived aldehydes with methoxy groups in enrichments (*15*). Overall, the cobalamin synthesis capability might be one of the key factors in explaining the ubiquitous distribution and predominance of genera *Baizomonas* and *Baizosediminiarchaeum* (previously subgroups-6/-5bb and -8), even the whole class Bathyarchaeia, in global freshwater and marine sediments (*25*).

### Atypical and flexible pathway for C1 compound metabolism

In addition to complex organics, Bathyarchaeia also exhibits striking potential in using multiple C1 compounds, including CO_2_, CO, formate, formaldehyde, methanol, methylamine and methane. For the orders Wuzhiqiibiales and Mazuousiales, the key genes of reductive glycine pathway (rGlyP), which is the seventh natural carbon fixation pathway reported recently (*46*), were identified in their MAGs, including the genes coding for glycine cleavage system (GCS), serine hydroxymethyltransferase (*glyA*) and serine deaminase (*sda*), as well as the H_4_-F methyl branch of WLP (Fig. 3a). Moreover, both analyzed Mazuousiales MAGs uniquely encode formate dehydrogenase (*fdh*), while the key genes (*coxM* and *coxS*) for aerobic CO dehydrogenase are largely found in the order Wuzhiqibiales (Fig. 4). These results suggest that some species from the orders Wuzhiqiibiales and Mazuousiales are likely able to assimilate CO_2_, CO and/or formate via the novel rGlyP. Distinct from the novel archaeal phylum Brockarchaeota that also has rGlyP (*47*), additional complete H_4_-MPT WLP and CO/formate metabolic potentials are identified in the orders Wuzhiqiibiales and Mazuousiales (Fig. 3 and 4), implying that these Bathyarchaeia lineages have more diversified and flexible capabilities in metabolizing different C1 compounds. Previous studies demonstrated that some Bathyarchaeia species could assimilate ^13^C-labeled carbonates, presumably through the H_4_-MPT WLP (*5, 14*). Here, we provide another possibility: a considerable number of Bathyarchaeia members are capable of autotrophic growth through the novel rGlyP. Given the prevalence of the orders Wuzhiqiibiales and Mazuousiales in modern marine sediments (*26, 41*), their overlooked metabolic capability and contribution to dark carbon fixation warrants further study.

**Fig. 4.**
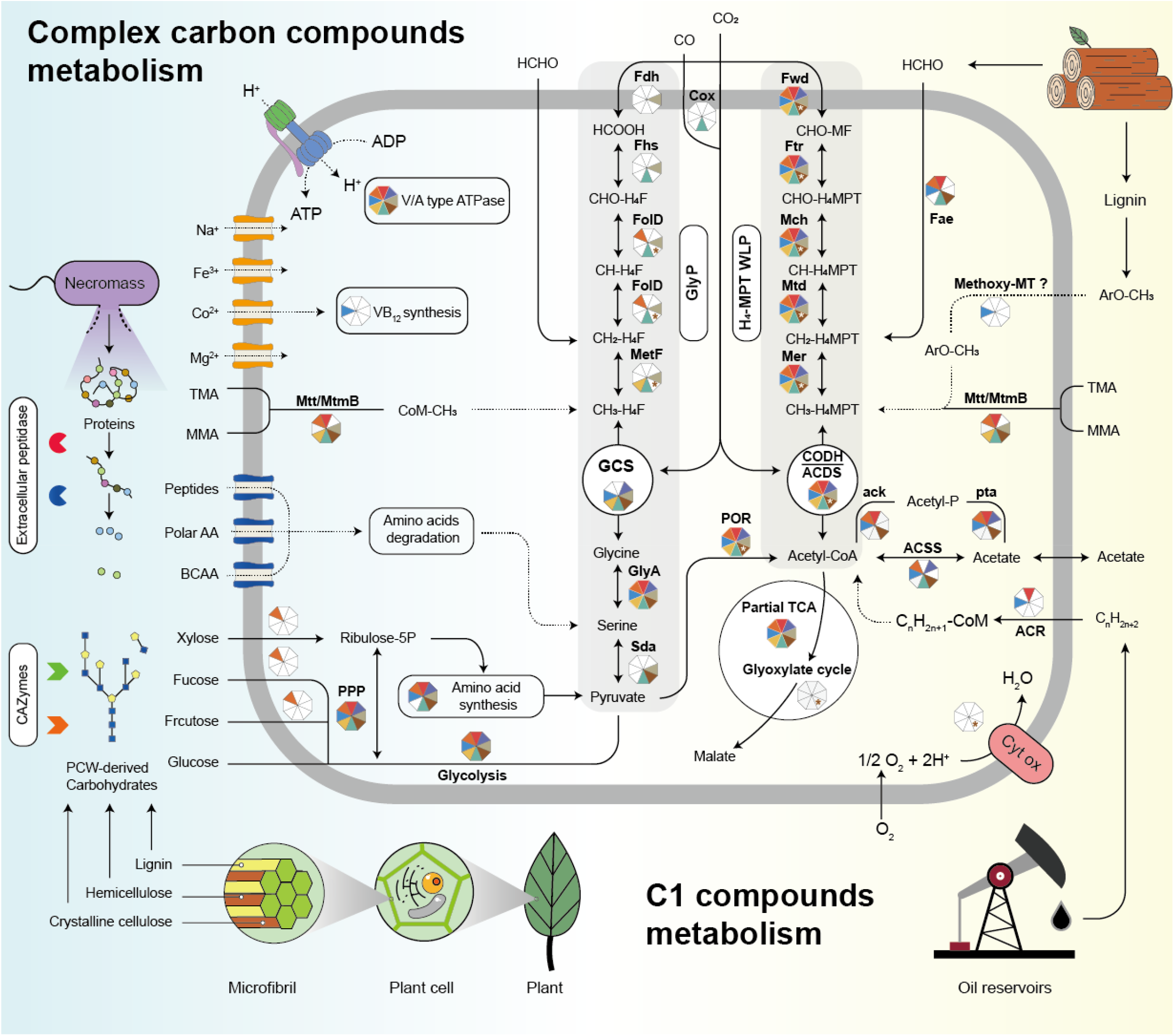
Overall metabolic reconstruction of the class Bathyarchaeia. Different pathways and gene presence for each Bathyarchaeia order are indicated by colored parts in octagon. White color denotes that the pathway or gene is absent in all MAGs of the order. The functional genes in the H_4_-MPT pathway marked with white stars and brown backgrounds, including Fwd, Ftr, Mch, Mtd, Mer and CODH/ACDS, indicate that they are present in all other MAGs in the order Houtuarculales, except for the special genus *Houtuousia* recovered from soil. In contrast, those genes marked with brown star in white background represent that they are exclusively identified in the members of genus *Houtuousia*, but not in other Houtuarculales MAGs. The following metabolic pathways and marker genes were used in the figure (see Table S6): PPP, archaeal pentose phosphate pathway; TCA cycle, tricarboxylic acid cycle; PFOR, pyruvate:ferredoxin oxidoreductase complex; ack-pta, acetate kinase and phosphate acetyltransferase; ACSS, acetyl-CoA synthetase; rGlyP, reductive glycine pathway; GCS, glycine cleavage system; GlyA, glycine hydroxymethyltransferase; SdaA, L-serine dehydratase; fae, formaldehyde-activating enzyme; Cox, aerobic carbon-monoxide dehydrogenase; Fwd, formylmethanofuran dehydrogenase complex; Ftr, formylmethanofuran – tetrahydromethanopterin N-formyltransferase; Mch, methenyltetrahydromethanopterin cyclohydrolase; Mtd, methylenetetrahydromethanopterin dehydrogenase; Mer, coenzyme F420-dependent N5,N10-methenyltetrahydromethanopterin reductase; Fdh, formate dehydrogenase; Fhs, formate-tetrahydrofolate ligase; *FolD*, methylenetetrahydrofolate dehydrogenase (NADP^+^)/methenyltetrahydrofolate cyclohydrolase; MetF, methylenetetrahydrofolate reductase (NADPH); Cyt ox, Cytochrome c oxidase; mttB, trimethylamine-corrinoid protein Comethyltransferase; mtmB, methylamine-corrinoid protein Comethyltransferase; ACR, alkyl-coenzyme M reductase complex; Compounds: TMA, trimethylamine; MMA, monomethylamine; BCAA, branched-chain amino acids; ArOCH_3_, methoxylated aromatic compounds.

The genes coding for mono- and trimethylamine-cobalamin methyltransferase (MT) systems (*mtmB* and *mttB*, respectively) are predicted in seven Bathyarchaeia orders (except for the sole MAG of the order Xuanwuarculales), most of which are distributed in the orders Wuzhiqibiales and Bifangarchaeales (Fig. 3a). Two Jinwuousiales MAGs also contain the encoding genes of methanol and dimethylamine-cobalamin methyltransferases (*mtaB* and *mtbB*, respectively). Under anoxic conditions, methylated C1 compounds are supposed to be mostly used by methanogens for methane production. However, it is unlikely for Bathyarchaeia, as the key gene coding for methyl-coenzyme M reductase (MCR) in methanogenesis is absent in the majority of Bathyarchaeia lineages (see the next part for details). In addition, anaerobic respiration and acetogenesis are alternative pathways for metabolizing methanol (*48*) and methylamines (*49*). Accordingly, given the generally missing sulfate reduction or denitrification pathways but the prevalent acetate metabolism-involved genes (e.g., Ack, Pta and ACSS) in most Bathyarchaeia MAGs (Fig. 3a), it is plausible that their MT system-encoded species are non-methanogenic methylotrophic archaea that could metabolize multiple methylated compounds via acetogenesis. Recently, ^13^C-labeled incubation experiments demonstrated active methylotrophy of methylamine and methanol by microbial communities from coalbeds 2 km below the seafloor with minor methanogenesis (*50*). Therefore, the non-methanogenic methylotrophic lifestyle might be an important adaptive strategy for microbial life in the extremely energy-limited deep biosphere, and so do these Bathyarchaeia that widely distributed in global deep marine sediments (*10, 29*).

Moreover, our other work also demonstrated that one *Baizosediminiarchaeum* species of the lignin-degrading clade employs a novel MT system to transfer the methyl groups from lignin-derived ArOCH_3_ to H_4_MPT, which are either further oxidized to CO_2_ or converted to acetate for energy production (*45*). This suggests that this Bathyarchaeia-specific MT system plays a key role in anaerobic lignin degradation, thus likely explaining their predominance in the lignin-enriched estuarine and nearshore sediments. In addition, formaldehyde is a common toxic C1 byproduct released during lignin-derived aromatic degradation (*51*). Here, the gene encoding formaldehyde-activating enzyme (*fae*) is widely identified in the analyzed MAGs, including the lignin-degrading clade, implying that formaldehyde could be detoxified or utilized via the H_4_-MPT WLP (*52*) for most Bathyarchaeia lineages.

Taken together, our results reveal the currently unrecognized and diverse roles of Bathyarchaeia in C1 compound transformation: a majority of them are inferred to be putative non-methanogenic methylotrophic archaea with the potential to metabolize different methylated compounds via diverse MT systems, including methanol, methylamines and ArOCH_3_. In comparison with Brockarchaeota (*47*), most of these Bathyarchaeia lineages possess an extra complete H_4_-MPT WLP and more methylamine-associated MT systems (Fig. 4), reflecting their more flexible methylotrophic strategy and substrate preference, the latter of which might facilitate their survival in the nutrient-limited deep subseafloor, as the amino residues released from methylamines are bioavailable nitrogen sources (*53*). Moreover, multiple methylated C1 compounds have been shown to be metabolized through methylotrophic methanogenesis and non-methanogenic methylotrophy in anoxic sulfate-reducing sediments (*54*), where Bathyarchaeia are also enriched (*55*). In conclusion, these results provide insights into the diverse and versatile C1 carbon metabolism of Bathyarchaeia by demonstrating their prevalent but overlooked methylotrophic capabilities and thus highlighting their unparalleled significance and contribution to the global carbon cycle.

Two MAGs (BA1 and BA2) from coalbed well formation water contain methyl-coenzyme M reductase (MCR)-encoding genes (*mcr*) (*16*), which were assigned to different families of the order Baizomonadales herein (Table S6). These *mcr*-like genes are inferred to be *acr* genes encoding novel alkyl-coenzyme M reductases (ACRs) involved in the anaerobic oxidation of multicarbon alkanes (*17, 56*). Interestingly, here we found a complete *mcr* gene cluster in another MAG (3300028193_21) from the order Jinwuousiales. These are the only three MAGs containing the *mcr* or *acr* genes in the 304 MAGs checked in this study, and as far as we are aware, also for all available Bathyarchaeia MAGs from public databases. Further phylogenetic analysis using *mcr*/*acrA* placed the two Baizomonadales-ACRs in the MCR group III, while the only MCR from the order Jinwuousiales was assigned to MCR group II (with mainly putative methanogens from the previous TACK superphylum and a few Archaeoglobi species) (Fig. S6). Previous studies indicate that the two *acr* genes of Baizomonadales might have been acquired via HGT events from other ACR-encoded euryarchaeotal hosts (*17, 57*). However, the Jinwuousiales *mcr* gene discovered here more likely evolved vertically from the common ancestor with other MCR-encoding archaea within MCR group II, because Bathyarchaeia and the most previous TACK members in this group, such as *Ca*. Methanomethylicia, *Ca*. Korarchaeia and *Ca*. Nezhaarchaeales, are reclassified to different class-or order-level units of the same phylum Thermoproteota in the GTDB taxonomy (*11*). Overall, such remarkable phylogenetic divergences among the three Bathyarchaeia *acr* genes not only reflect their complex evolutionary history of methane metabolism but also expand their putative ecological roles in alkane cycling (methanogenesis or alkane oxidization).

### Co-evolution of Bathyarchaeia and the early Earth surface environment

By using the temporal constraints of an HGT event of the chromosome segregation (SMC) gene (*58*), the common ancestor of Bathyarchaeia is predicted at ∼3.37 Ga (posterior 95% confidence interval (CI): 3.62-3.13 Ga), coinciding with the subaerial emergence of the first continent on Earth ∼3.46 to 3.2 billion years ago (*59, 60*) (Fig. 5 and Fig. S7). In the absence of plant root systems, Paleoarchean surface terrains would have been subject to rapid migration of riverbeds, and given that the tidal range was larger in the past, it is likely that emergent continental crust would have experienced enhanced wetting (*61*). Therefore, the ancient terrestrial phototrophic microbial communities, in particular cyanobacteria, would colonize the exposed land surface and grow as biological soil crusts and microbial mats (62). These mats certainly could have been sloughed off and transported by rivers to coastal settings where their detritus accumulated in the sediment pile. Concomitant with increased continental weathering, the enhanced transport of suspended and dissolved loads to the oceans would cause the ultimate accumulation of sediments to form shallow continental shelves with increased nutrient supply (e.g., phosphorus). As discussed above, modern Bathyarchaeia species have versatile heterotrophic capabilities and mostly inhabit anoxic coastal sediments, which are characterized by rapid terrestrial sedimentation rates. Accordingly, it is reasonable to infer that the ancestral Bathyarchaeia originated in these newly formed coastal settings associated with continental emergence on the Archean Earth. More importantly, the presence of heterotrophic Bathyarchaeia so early in Earth’s history not only implies abundant organic carbon availability but also points to an early terrestrial biosphere that directly arose through the creation of habitable space on land.

**Fig. 5.**
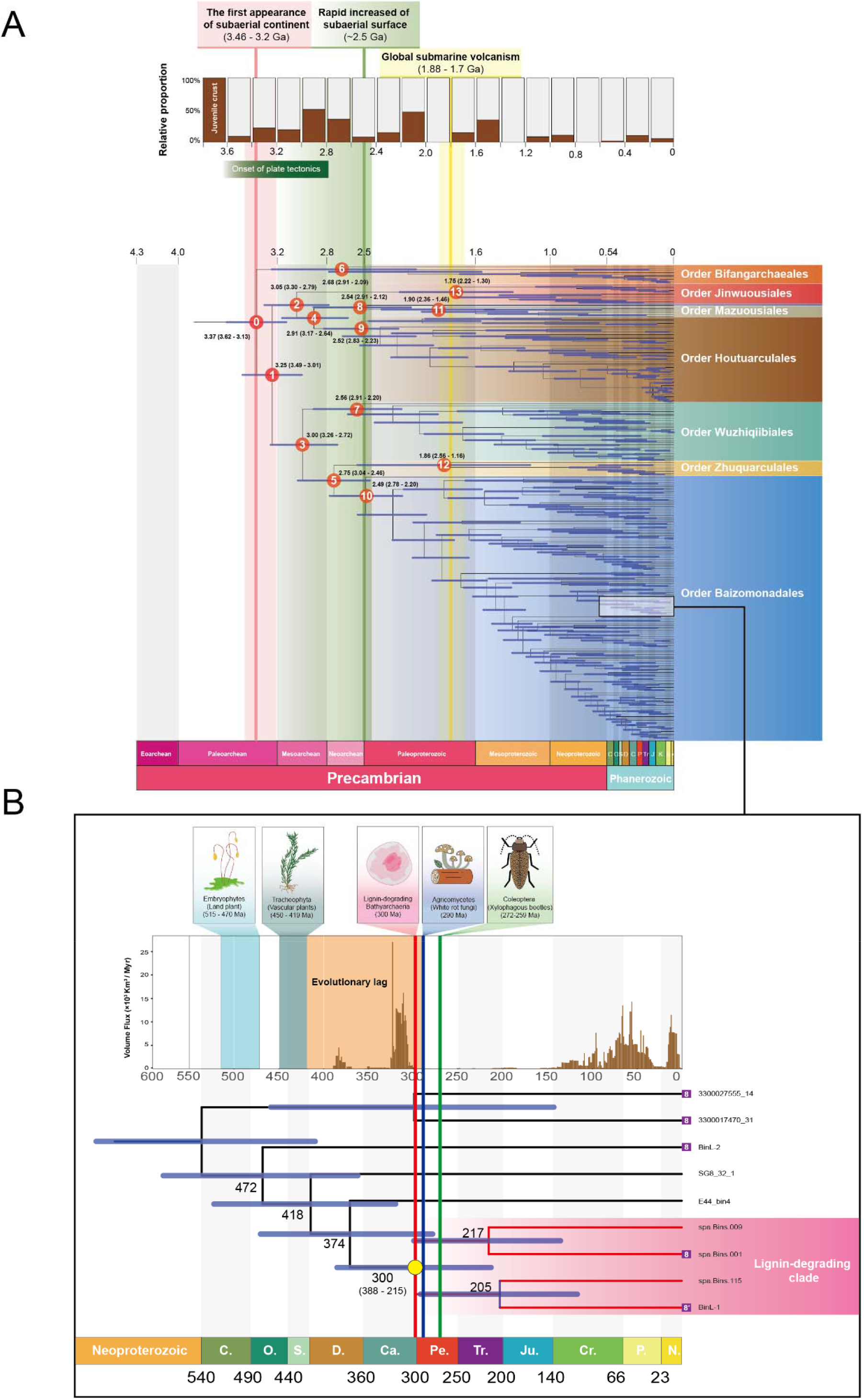
Evolutionary history of the class Bathyarchaeia and timing correlations with major geological activities. **A**, The phylogenomic tree and estimated divergence times of the major Bathyarchaeia lineages. The whole tree was constructed based on 259 Bathyarchaeia and 190 reference MAGs by the concatenated alignment of their SMC and 16 conserved proteins using the best-fit model *LG+F+R10+C60* in IQ-TREE2 (see Fig. S7). Molecular dating was calculated by MCMCtree with three different calibration time points, and the timeframe displayed here is based on the calibration priors > 1.7 Ga for the lineage of Nostocales, 2.5-3.0 Ga for the crown group of oxygenic Cyanobacteria and estimated divergence times of Archaea and Bacteria at 3.8-4.29 Ga. Different color schemes represent the descendent lineages for eight Bathyarchaeia orders (the label of order Xuanwuarculales is removed as the only MAG). The divergence ages of major nodes are numbered 0-13 and labeled with the posterior 95% confidence intervals (flanking horizontal blue bar). The relative proportions of juvenile crust through time are represented as a brown bar chart in the top panel, which is deduced from the Nd model age of sedimentary rocks (*72*). The vertical pink band and line indicate the estimated period (3.46-3.2 Ga) of the subaerial continent first exposed on the early Earth (*60, 61*) and the corresponding origin time of the common ancestor of Bathyarchaeia ∼3.37 Ga, respectively. The green band represents the time interval from the emergence of the first subaerial continents to the rapidly increasing subaerial landmass at ∼2.5 Ga (the green line) (*75*). The yellow band shows the timing of global submarine volcanism and hydrothermal activity intensively occurred at 1.88-1.7 Ga (*77, 78*). **B**, The evolutionary timeline of the Bathyarchaeia lignin-degrading lineage and contemporaneous Permo-Carboniferous coal peak. The tree is part of the whole phylogenomic tree of Bathyarchaeia, comprising nine MAGs from the genus *Baizosediminiarchaeum* within the order Baizomonadales. The clade with a pink background has four MAGs (spa.Bins.001, spa.Bins.115, spa.Bins.009 and BinL-1), which were specifically recovered from the lignin enrichment and designated the Bathyarchaeia lignin-degrading clade in this study. The MAGs labeled with the number “8” within the purple box are assigned to the previous subgroup-8. The yellow dot indicates the estimated origin timing of the common ancestor for this lignin-degrading clade at ∼300 Ma (CI: 388-215 Ma). The top panel shows the burial flux of terrestrial organic sediments accumulated in North America through time (*80*) and contemporaneous key evolutionary time points for terrestrial plants (*124, 125*), white rot fungi (*83*) and xylophagous beetles (*86, 87*). The red, blue and green lines represent the estimated origin timing (period) of the Bathyarchaeia lignin-degrading clade, white rot fungi and xylophagous beetles, respectively.

The Paleoarchean rock record certainly points to the presence of stromatolites that grew in the shallow waters fringing the newly exposed continental landmass. For example, the ∼3.42 Ga Buck Reef Chert contains fine anastomizing carbonaceous laminations that have been interpreted as fossil microbial mats (*63*). Similarly, one of the oldest known stromatolites, the ∼3.43 Ga Strelley Pool Formation in northwestern Australia, contains complex morphologies that vary along a carbonate platform that is several kilometers long and has been interpreted as a microbial reef (*64, 65*). More recently, the earliest biosignatures were discovered from the ∼3.5 Ga hot spring deposits (*66*). What is consistent among these deposits is the influence of hydrothermal activity in association with frequent volcanic eruptions.

Interestingly, we found that the orders Bifangarchaeales and Jinwuousiales are specific thermophilic lineages inhabiting hot spring and hydrothermal sediments, respectively (Fig. 2b). As the two most ancient lineages, phylogenetic results reveal that the common ancestor of order Bifangarchaeales diverged from the root directly, followed by the emergence of order Jinwuousiales at ∼3.05 Ga (CI 3.30-2.79 Ga) (Fig. 5 and Table S8). This implies that Bathyarchaeia very likely evolved in an ancient hydrothermal setting, in accordance with the hot origin theory proposed before (*28*). In this context, we inferred that the common ancestor of Bathyarchaeia might emerge from the shallow submarine volcanic (or terrestrial hot spring) sediment enriched with organics from either continental weathering or *in situ* production by microbial mats. In addition, the early Earth surface environment is likely to have accreted large amounts of prebiotic methylated compounds from cometary delivery (*67*). Together with the enriched MT systems identified in the two ancient thermophilic orders (Fig. 3a), we suggest that the ancestral Bathyarchaeia might be non-methanogenic methylotrophic archaea, and this view is consistent with a recent study suggesting the methylotrophic origin of methanogenesis at ∼3.8-4.1 Ga ago (*20*).

The first extensive diversification of Bathyarchaeia occurred near ∼3.0 Ga, including monophyletic divergences of the orders Jinwuousiales and Wuzhiqiibiales from other Bathyarchaeia lineages (Fig. 5a). Although the specific timing and mechanism remain highly controversial (*68*), ∼3.0 Ga has been interpreted as a pivotal period during the evolution of Earth’s tectonics (*69*–*71*). At that time, the majority (∼65%) of juvenile continental crust had already formed (*71*), along with dramatically altered chemical composition (*70*), increased reworking/growth rate (*69, 72*) and thickness (*71*), in turn marking the putative onset of plate tectonics (*73*). Indeed, this timing is coincident with the Kaapvaal and Pilbara cratons (*74*). As the cratons were in isostatic uplift, the felsic rocks became exposed to erosion and weathering, leading to enhanced export of nutrients to the Archaean ocean. We suggest that increased weathering led to higher nutrient delivery, primary productivity and organic carbon burial, which drove the evolution of certain Bathyarchaeia lineages, such as peptides-utilizing Wuzhiqiibiales, to adapt to the great change in sediment composition.

The second extensive diversification of Bathyarchaeia occurred between ∼2.75 and ∼2.49 Ga, during which the taxonomic radiations of the orders Wuzhiqiibiales (∼2.56 Ga), Houtuarculales (∼2.52 Ga) and Baizomonadales (∼2.49 Ga), as well as the emerged common ancestor of Mazuousiales and Xuanwuarculales (∼2.54 Ga) (Fig. 5a and Table S8). This timing is associated with the rapid and expansive emergence of the subaerial landmass at the Archean-Proterozoic boundary (*75*), similar to the ∼3.0 Ga, which significantly increased subaerial weathering (*76*), the burial rate of organic carbon and the extent of continental and sediment volume (*62*).

Interestingly, the phylogenetic radiation of different Bathyarchaeia lineages with diversified carbon metabolism might have been triggered by changes in emergent crustal lithology. For instance, between ∼3.5 and ∼3.2 Ga, the crust was primarily mafic, but by ∼3.0 Ga, it became more felsic in composition (*70*). Additionally, as a consequence of adapting to the cooler Earth surface condition, the depositional environments for Bathyarchaeia changed from inhabiting early geothermal environments and then transitioning to moderate or cold modern marine sediments. Along those lines, the third major diversification event, leading to taxonomic radiation of two hydrothermal orders Jinwuousiales and Zhuquarculales, occurred at ∼1.86 (CI: 2.56-1.16) and ∼1.75 (CI: 2.22-1.30) Ga, respectively. This time window was concordant with a short period of intense submarine volcanic activity that took place globally at ∼1.88-1.7 Ga (*77, 78*). Therefore, the geological consequences of rapid continental growth and global magmatic eruption approximately 1.8 billion years ago might have significantly impacted the interior diversification of the ancestral hydrothermal Bathyarchaeia, probably by isolating them into distinct ecological niches.

### Lignin-degrading Bathyarchaeia clade potentially contributed to the end of Permo-Carboniferous coal production peak

An unparalleled interval of carbon sequestration occurred during the late Carboniferous to early Permian (∼323–252 Ma), which corresponded to the greatest coal deposits in Earth’s history (*79, 80*), the so-called Permo-Carboniferous coal production peak. As the second most abundant biopolymer on Earth, lignin is sourced from dry woody (vascular) plant material. It is extraordinarily resistant to biological degradation due to its highly amorphous and polyphenolic structure (*81*), making it the main precursor of coal. Today, white-rot fungi are the primary lignin degraders, except for a few bacterial species with very limited capabilities for delignification (*81*). Thus, Robinson et al. (*82*) hypothesized that a temporal gap of ∼120 Myr between the origin of lignin-synthesizing vascular plants and the delayed evolution of lignin-degrading white rot fungi provided a window of opportunity for a substantial amount of recalcitrant vegetal polymers to accumulate and, in turn, become coal. The subsequent molecular dating evidence makes this paradigm more convincing by demonstrating that the common ancestor of white rot fungi did not evolve until ∼290 Ma (*83*). Recently, certain Bathyarchaeia species from the genus *Baizosediminiarchaeum* (previous subgroup-8) have been reported to be capable of lignin utilization under anaerobic conditions (*14*). Here, four MAGs of the genus *Baizosediminiarchaeum* that were specifically recovered from the lignin-enrichment culture in our lab were designated as the lignin-degrading Bathyarchaeia clade.

To increase the resolution of the phylogenetic signal of its adjacent lineages, an additional 21 single conserved genes were added to the phylogenetic marker set for the molecular dating of this lignin-degrading clade (see Materials and Methods for details). Phylogenetic analysis suggests that the lignin-degrading lineage is monophyletically evolved from one common ancestor, whose divergence time was estimated at ∼300 Ma (CI: 388-215 Ma), coinciding with the emergence of ancestral white rot fungi and the peak of carbon sequestration at the end of the Carboniferous period (Fig. 5b). Therefore, we infer that the lignin-degrading Bathyarchaeia ancestor also likely contributed to the sharp decline in Permo-Carboniferous coal production, similar to what white rot fungi did according to the evolution lag theory.

More importantly, distinct from the aerobic white rot fungi, this special Bathyarchaeia clade can degrade lignin under strictly anoxic condition, which means that their ancestor could have continuously consumed buried woody material over hundreds of million years during the coal formation process (*79, 84*). Thus, they played a much more important role in determining organic carbon sequestration than their fungal counterparts, which are strictly restricted in the subaerial oxic environment. Furthermore, the putative existence of anaerobic lignin-degrading archaea 300 million years ago makes anoxic conditions no longer a vital factor determining the fate of recalcitrant woody material after burial in Carboniferous sediments. This finding directly challenges the alternative abiotic hypothesis that explains the unusual coal abundance as a confluence of the disappearance of widespread anoxic waterlogged environments driven by dramatic climate and tectonic changes (*80, 85*). Moreover, the origin of xylophagous beetles (order Coleoptera) was also coeval with the two microbial lignin degraders (*86, 87*) (Fig. 5b). Given that their modern descendants accounted for ∼29% of the deadwood degradation in modern forest ecosystems (*88*), the contemporaneous ancestral xylophagous insects might also have contributed to the decline of the Permo-Carboniferous coal peak. Taken together, our results provide supporting evidence for the evolution lag theory in explaining the unusual coal abundance during the Permo-Carboniferous period. Most importantly, the improved evolutionary lag theory is an excellent case for illustrating the feedback and co-evolution between life and Earth over geological timescales, in which Bathyarchaeia, as the most abundant and widespread microorganism on Earth today, might have played an incomparable role in global carbon cycling at least ∼3 million years ago.

In summary, this study provides a comprehensive vision of the phylogeny, metabolism and evolution of Bathyarchaeia within an improved phylogenetically congruent taxonomic framework, which is established based on the largest Bathyarchaeia MAG dataset and GTDB methodology to date. Comparative genomics results show the distinct environmental distribution and diversified carbon metabolism among different Bathyarchaeia lineages. In particular, multiple atypical C1 metabolic pathways, including different methyltransferase systems, the reductive glycine pathway and the newly discovered methyl-coenzyme M reductase, reflect their previously overlooked potential roles as anaerobic methylotrophs, dark carbon fixers and methane/alkane oxidizers. Moreover, the evolutionary history of Bathyarchaeia is linked to tectonic evolution on early Earth, indicating that the emergence of the first continent might have triggered the origin of Bathyarchaeia from an ancient geothermal sedimentary environment, while the subsequent continental expansion and intensive hydrothermal activity probably facilitated the dramatic diversifications of different Bathyarchaeia lineages. Interestingly, the temporal coincidence between the appearance of the lignin-degrading Bathyarchaeia clade and white rot fungi implies that certain ancient Bathyarchaeia groups likely contributed to the sharp decline in coal deposits during the Permo-Carboniferous period.

Our work provides systematic insights into the taxonomic and metabolic diversity of the whole class Bathyarchaeia, which not only leads to a better understanding of their overall ecological and biogeochemical significance, but also lays the foundation for future ecological and functional studies of different Bathyarchaeia lineages. This study provides a good case for illustrating the feedbacks and co-evolution between life and Earth, which further expands the roles of Bathyarchaeia in carbon cycling from modern Earth to the whole geological timescale. Nevertheless, future wider ecological investigations and continuous efforts in cultivating or isolating pure strains are needed to further decipher their specific roles in modern environments and potential implications in deep time.

### Nomenclature and Etymology

#### Description of genus *Bifangarchaeum* gen. nov

*Bifangarchaeum* (N.L. masc. n. Bi Fang, A single-legged bird in Chinese mythology, usually regarded as the symbol of wildfire; N.L. neut. n. archaeum, an archaeon; *Bifangarchaeum*, the archaeon associated with terrestrial hot environments. The type genus of family *Ca*. Bifangarchaeaceae and order *Ca*. Bifangarchaeales). The nomenclatural type of this genus is *Bifangarchaeales caldum*^Ts^.

#### Description of genus *Jinwuousia* gen. nov

*Jinwuousia* (N.L masc. n. Jin Wu, A three-legged bird living on the Sun in Chinese mythology; L. fem. n. ousia, an essence; *Jinwuousia*: a microbe living in hot environments. The type genus of family *Ca*. Jinwuousiaceae and order *Ca*. Jinwuousiales). The nomenclatural type of this genus is *Jinwuousisles tripozesto*^Ts^.

#### Description of genus *Houtuarcula* gen. nov

*Houtuarcula* (N.L masc. n. Hou Tu, Chinese earth god; L. fem. n. arcular, small box; *Houtuarcula*: a terrestrial archaeon. The type genus of family *Ca*. Houtuarculaceae and order *Ca*. Houtuarculales). The nomenclatural type of this genus is *Houtuarcula pratum*^Ts^.

#### Description of genus *Mazuousia* gen. nov

*Mazuousia* (N.L fem. n. Mazu, Chinese sea goddess; L. fem. n. ousia, an essence; *Mazuousia*: a microbe inhabiting the oceanic environments. The type genus of family *Ca*. Mazuousiaecae and order *Ca*. Mazuousiales). The nomenclatural type of this genus is *Mazuousia monanthrakas*^Ts^.

#### Description of genus *Xuanwuarcula* gen. nov

*Xuanwuarcula* (N.L masc. n. Xuan Wu, A black turtle with snake tail, known as the God of North and symbol of iciness and water in Chinese mythology; L. fem. n. arcular, small box; *Xuanwuarcula*: the archaeon living in cold underground water. The type genus of family *Ca*. Xuanwuarculaceae and order *Ca*. Xuanwuarculales). The nomenclatural type of this genus is *Xuanwuarcula inferioraqua*^Ts^.

#### Description of genus *Wuzhiqiibium* gen. nov

*Wuzhiqiibium* (N.L masc. n. Wu Zhiqi, God of Huai river in Chinese mythology, the prototype of Sun Wukong (the legendary Chinese figure “Monkey King”); L. neut. n. bium, bios, life; *Wuzhiqiibium*: the microbes associated with estuarine environments. The type genus of family *Ca*. Wuzhiqiibiaceae and order *Ca*. Wuzhiqiibiales). The nomenclatural type of this genus is *Wuzhiqiibium servoceani*^Ts^.

#### Description of genus *Zhuquarcula* gen. nov

*Zhuquarcula* (N.L masc. n. Zhu Que, A red bird as the God of South and symbol of fire in Chinese mythology; L. fem. n. arcular, small box; *Zhuquarcula*: the archaeon living in hot environments. The type genus of family *Ca*. Zhuquarculaceae and order *Ca*. Zhuquarculales). The nomenclatural type of this genus is *Zhuquarcula teneretur*^Ts^.

#### Description of genus *Baizomonas* gen. nov

*Baizomonas* (N.L masc. n. Bai Ze, A propitious white creature living in the cold Kunlun Mountains in Chinese mythology; L. fem. n. monas, a monad; *Baizomonas*: the microbe associated with cold environments. The type genus of family *Ca*. Baizomonadaceae and order *Ca*. Baizomonadales). The nomenclatural type of this genus is *Baizomonas nivis*^Ts^.

## Method and Materials

### Sample collection and DNA extraction

In this study, two natural sediment samples and four sediment incubation enrichment cultures were sequenced by our lab and utilized to achieve more Bathyarchaeia MAGs. All necessary information involved in sampling and cultivation conditions is listed in Table S1.

#### Deep sea sediment from South China Sea

One 8.3 m piston core named QDN-14B was sampled from the Haima cold seep located in the South China Sea during the cruise ‘20150402’ R/V Haiyang □ in April 2015, and the specific subsampling and preservation method was described in a previous study (*89*).

#### Coastal sediment from East China Sea

One 6 m gravity core A2-3 was collected from the Yangzi estuary (30.927933°N, 122.473233°E) in the East China Sea during a cruise in July 2017. The subsamples used for molecular analysis were cut into 10-cm intervals, kept in liquid nitrogen during transport to the laboratory, and stored at -80 □ until further analysis.

#### Sediment enrichment with lignin

Four sets of sediments from different natural environments were aerobically incubated with lignin to enrich Bathyarchaeia. One intertidal sediment sample (DYS) was collected from Dayangshan Island (30.592817°N, 122.083493°E), Hangzhou Bay, China, as described previously (*14*). One freshwater sediment sample (PYH) was obtained from Poyang Lake, Jiangxi, China. The other two sediment samples (SQ and SRBZ) were collected from two hot spring pools with different temperatures in Tengchong, China. The detailed cultivation method for the DYS set was described (*14*), and the cultivation conditions for the other three sets are listed in Table S1.

For enrichment cultures, the DNA extraction method was the same as previously described (*14*). A modified SDS-based method (*90*) was utilized to extract high-quality DNA from two natural sediment samples for metagenomic library construction and sequencing.

### Metagenomic sequencing, assembly and binning

For all sediment samples mentioned above, metagenomic libraries were constructed and sequenced on the HiSeq X Ten platform (2 × 150 bp paired-end reads). First, raw metagenomic reads for each sample were trimmed by using Trimmomatic v0.39 (*91*) with default parameters and possible adaptors. After quantity control, clean reads were assembled into contigs by using MEGAHIT v1.2.9 (*92*) with the parameter *-min-count 2 -k-min 41 -kmin-1pass -k-max 147 -k-step 10*, followed by mapping clean reads to contigs by using bowtie2 v2.4.4 (*93*) with a *-very-sensitive* pattern for estimating the sequencing coverage of each contig.

Metagenomic binning was performed individually for each sample by a modified protocol from85. First, MetaBAT2 v2.2.15 (*94*) and MaxBin v2.2.7 (*95*) were utilized to recover MAGs with contigs more than 1k (1.5k for MetaBat2), 3k and 5k in each metagenomic sample, respectively. The resulting MAGs were refined by removing contigs with incongruent taxonomic information and divergent genomic properties (e.g., GC content and coverage) based on the “outliers” pattern from RefineM v0.0.22 (*96*). All refined MAGs from the same sample were integrated into optimized, non-redundant bins by using DAS Tool 1.1.4 (*97*). The quality and initial taxonomic classification of integrated MAGs were estimated by checkM (*98*) and GTDB-Tk (*23*), respectively. High-quality MAGs (completeness > 90% and contamination < 10%) assigned to Bathyarchaeia were retained for downstream analysis.

### Public data collection

Candidate Bathyarchaeia MAGs were downloaded from the genome reference databases of NCBI (https://www.ncbi.nlm.nih.gov/), GTDB (Genome Taxonomy Database) (*11*) and GEM (Genome from Earth’s Microbiome) catalog (*22*) based on the key words “Bathyarchaeota,” “MCG” (miscellaneous Crenarchaeota group) or “Bathyarchaeia” (final data collection in December 2020). The quality of these candidate Bathyarchaeia MAGs was estimated by checkM (*98*). In addition, sampling and environmental information for each available candidate, Bathyarchaeia, were manually searched and curated in the corresponding literatures.

### MAG refinement and subgroup affiliation

Due to the overlapping among databases and possible incorrect taxonomic assignment, all available candidate Bathyarchaeia MAGs from public databases and our own datasets were carefully checked for potential false positivity in taxonomic classification as “Bathyarchaeia” by using the *classify_wf* function in GTDB-Tk (*23*). After removing the MAGs assigned to other microbial lineages, the average nuclear acid identities (ANIs) between pairwise sets of MAGs were calculated by using FastANI (*99*). The taxonomic level for species was defined with an ANI threshold higher than 95% (*100*). The MAGs were clustered into different groups if their ANIs among each other were higher than the species threshold. The MAGs with the best quality score (completeness – contamination × 5) were selected as the representative MAG of this group and retained for downstream analysis. In general, the genome quality of selected representative Bathyarchaeia MAGs satisfy the median quality based on the MIMAG standard, a few exceptions exist in order to retain some well-known or important Bathyarchaeia species, such as the first single-cell sequenced Bathyarchaeia genome (SCGC AB-539-E09) (*3*), which facilitate the further discussion about their metabolic characteristics and evolutionary history. The accession number and source for each representative Bathyarchaeia MAG are listed in Table S2.

For each representative Bathyarchaeia MAG, their 16S rRNA genes were initially identified and extracted by using Barrnap (https://github.com/tseemann/barrnap) and manually curated BLASTN check in SILVA 138.1 SSU Ref database for potential contamination (*101*). The reference dataset includes two parts: the first is the 16S rRNA genes with confirmed subgroup classification information from Zhou et al. (*10*), while the other is collected from SILVA 138.1 SSU Ref database. The reference sequences from SILVA were clustered by using CD-HIT (https://github.com/weizhongli/cdhit) with 90% identity and compared with the first dataset for de-redundancy. Subsequently, the non-redundant reference dataset, 16S rRNA genes identified from Bathyarchaeia MAGs and additional archaeal outgroup sequences were aligned by using SILVAngs (https://ngs.arb-silva.de/silvangs) and trimmed by trimAI with the *-automated1* pattern (*102*). The maximum likelihood phylogenetic tree was constructed by using raxML version 8.2.12 (*103*) with the *-m GTRGAMMA -N autoMRE* parameter. Note that some representative Bathyarchaeia MAGs without 16S rRNA genes were assigned to subgroups identical to their co-species (ANI > 95%) from the same cluster if the latter had the subgroup-assigned 16S rRNA gene (Table S3).

### Phylogenetic rank normalization and nomenclature

*Phylogenomic inference*. A total of 304 representative Bathyarchaeia MAGs were selected to construct the phylogenomic tree for taxonomic rank normalization, with an additional 55 archaeal genomes from other phyla (and other orders from the phylum Thermoproteota) as the reference dataset (Table S2). Their 122 archaeal-specific conserved proteins were identified, aligned, trimmed individually and concatenated together by using the identity function in GTDB-Tk (*23*). The maximum likelihood phylogenomic tree for these 359 archaeal MAGs/genomes was inferred by IQ-TREE v2.1.2 (*104*) using the best-fit *LG+F+R10+C60* model with 1000 ultrafast bootstrap replicates and 1000 bootstrap replicates for SH-aLRT and visualized on the iTOL website (*105*).

#### Rank classification and normalization

The taxonomic rank for each Bathyarchaeia MAG was initially inferred by using the *classify_wf* function in GTDB-Tk (*23*) and further curated carefully in consideration of the phylogenetic topological structure and environmental information of each lineage. The modified taxonomic ranks of Bathyarchaeia were normalized and evaluated by PhyloRank v0.1.10 (https://github.com/dparks1134/PhyloRank) based on the relative evolutionary divergence (RED) values of each taxonomic rank (Table S4).

#### Automated nomenclature

Given the difficulty in manually creating hundreds of new genus names for Bathyarchaeia MAGs, an automated approach was applied to solve this problem by GAN (*24*) based on three individual word roots from traditional Chinese mythological figures, environmental terms, and universal Latin endings or diminutives provided by ICNP (International Code of Nomenclature of Prokaryotes) (Table S9). For each proposed genus, their own type materials are designated as those with genomic quality that meets the defined standard (*106*) (or only the best one in all MAGs if no other MAGs satisfy the standard) and environmental representativity, which are utilized to define the names of the genus and corresponding higher-level taxonomic ranks by adding an appropriate suffix to the stem based on the ICNP rules (*107*). The nomenclature and etymology of each taxonomic rank are listed in Table S5.

### Genomic analysis and functional annotation

#### Genomic characteristics

General genomic traits, including genome size, GC content, predicted gene number, and coding density, were accessed from the checkM results directly. C-ARSC (carbon atoms per amino acid residue side chain) and N-ARSC (nitrogen atoms per amino acid residue side chain) were calculated by using custom scripts from Feng et al. (*108*). The significant differences in each genomic characteristic among different orders were compared by using the Wilcoxon test with Bonferroni correction (Fig. S3).

#### Functional gene annotation

To minimize the error possibility from incompleteness or contamination, 86 high-quality MAGs were selected as representative Bathyarchaeia MAGs to perform metabolic reconstruction in detail. ORFs (open reading frame) were predicted from each MAG by Prodigal V2.6.3 (*109*) in *single* mode and further annotated in a series of databases in parallel, including KEGG (*110*), Pfam (*111*), and UniProt (*112*), by using DRAM (*113*) with default parameters. The resulting annotation for specific key genes was further verified across at least two databases or manually checked based on the self-built custom database using Diamond blastp v0.9.14 (*114*). Genes encoding carbohydrate degradation enzymes described in the carbohydrate-active enzymes (CAZYmes) database V10 (*115*) were searched by HMMER 3.3.2 (*116*) with a threshold e-value < 1e^-15^, while peptidase and proteinase genes were detected in the MEROPS database 12.4 (*117*) using Diamond blastp v0.9.14 (*114*) with a threshold of coverage > 30% and e-value < 1e^-10^. The functional gene annotation of each Bathyarchaeia MAG is listed in Table S6.

#### Phylogenetic analysis of McrA

The phylogenies of McrA protein sequences retrieved from the representative Bathyarchaeia MAGs were further analyzed in this study. The phylogenetic tree of McrA was constructed together with the reference sequences from a previous study on anaerobic alkane-oxidizing archaea (*17*) by IQ-TREE v2.1.2 (*104*) with the best-fit *LG+F+G+C60* model.

### Phylogenetic divergence time estimation

#### Phylogenomic analysis

The molecular dating analysis of Bathyarchaeia uses the same methodology as previous studies on the evolution of methanogens (*59*) and methanogenesis (*20*), which are based on a temporal constraint of horizontal gene transfer (HGT) events of the structural maintenance of chromosome (SMC) proteins from certain euryarchaeotal methanogen lineages to the ancestor of Cyanobacteria (*59*), for which no unambiguous fossil records are available. For this purpose, a total of 259 high- and median-quality Bathyarchaeia MAGs were selected for phylogenetic analysis, together with an additional 190 reference genomes/MAGs from representative Cyanobacteria and other archaeal phyla (Table S10). Sixteen single copy conserved proteins and the structural maintenance of chromosomes (SMC) protein were predicted from each analyzed MAG after careful curation. To check the phylogenetic conservation and potential HGT events of each marker gene, their encoding protein sequences were aligned with MAFFT v7.310 (*118*), and trees were constructed individually by iQ-TREE2 with their own best-fit models (Table S11). The results support that they are conserved phylogenetic marker genes for the class Bathyarchaeia and suitable for the next molecular dating analysis (Fig. S8). The protein alignments of 16 conserved marker genes and the SMC gene were concatenated with FASconCAT v1.0 (*119*) and analyzed by IQ-TREE v2.1.2 (*104*) under the best-fit *LG+F+R10+C60* model with 1000 ultrafast bootstrap replicates and 1000 bootstrap replicates for SH-aLRT.

#### Divergence time estimation

The molecular dating analysis was conducted by using MCMCtree in paml version 4.9j with the *WAG* model (*120*) and iteratively run until convergence. Three nodes were calibrated on the phylogenomic tree: (1) The predicted differentiation time of bacterial and archaeal lineages at ∼4.29-3.8 Ga (*121*). (2) The divergence time of Cyanobacteria is estimated at 2.5-3.0 Ga (*122*). (3) Fossil evidence suggested that cyanobacterial clades Nostocales and Stigonematales already exist at ∼2.0 to 1.2 Ga (*59, 123*), which was defined as >1.2, >1.7 and >2.0 Ga, respectively, and calculated in the next divergence time estimation individually (*21*).

#### Alternative dating strategy for the lignin-degrading clade

Although the phylogenetic topology of the whole class Bathyarchaeia is rather robust based on the marker set of 16 single copy conserved proteins plus SMC protein, the phylogenies of the Bathyarchaeia lignin-degrading clade and close lineages show slight instability, which is attributed to the absence of marker genes in their MAGs (Table S12). To address this issue, another 21 single conserved marker genes were selected to supplement the marker gene set to provide enough phylogenetic information for this special clade (Table S12). In this way, a total of 37 marker genes coding for proteins plus the SMC protein were utilized to construct the phylogenomic tree of the whole Bathyarchaeia lineage for estimating the divergence time of the lignin-degrading clade. The dating method and calibration time points are exactly the same as described before.

## Acknowledgments

We are thankful to Dr. Philip Hugenholtz for his valuable suggestions in GTDB classification and nomenclature and to Sidra Ishaq for her advice in Etymology. We thank Chen Li, Yujue Wang, and Ya Guo from SJTU HPC for useful discussion of software. We are grateful to the researchers who shared their sequence data on the NCBI (https://www.ncbi.nlm.nih.gov/) and to the US Department of Energy Joint Genome Institute (http://www.jgi.doe.gov/) for providing genome files in collaboration with the scientific community.

## Author contributions

- Conceptualization: JLH, FPW, YZW
- Methodology: JLH, YZW
- Investigation: PFZ, NY, TTY, LLW and MYN
- Visualization: JLH
- Supervision: FPW and YZW
- Writing—original draft: JLH
- Writing—review & editing: JLH, FPW, YZW, KK

## Data and materials availability

The MAGs generated by our lab in the current study have been submitted under NCBI BioProject accession number PRJNA869847 and eLMSG (an eLibrary of Microbial Systematics and Genomics, https://www.biosino.org/elmsg/index) under accession numbers LMSG_G000011401.1-LMSG_G000011440.1. All MAGs from the GEM (Genome from Earth’s Microbiome) catalog are publicly available at https://genome.jgi.doe.gov/portal/GEMs/GEMs.home.html. The specific accession number for each MAG analyzed in this study is provided in Table S2.

