## Supplementary Materials for "Taxonomic and carbon metabolic diversification of Bathyarchaeia during its co-evolution history with the early Earth surface environment"

Science Advances

AAAS

Jialin Hou *et al.*

**This PDF file includes:**

Figs. S1 to S8
Legends for tables S1 to S12

**Other Supplementary Materials for this manuscript include the following:**

Tables S1 to S12

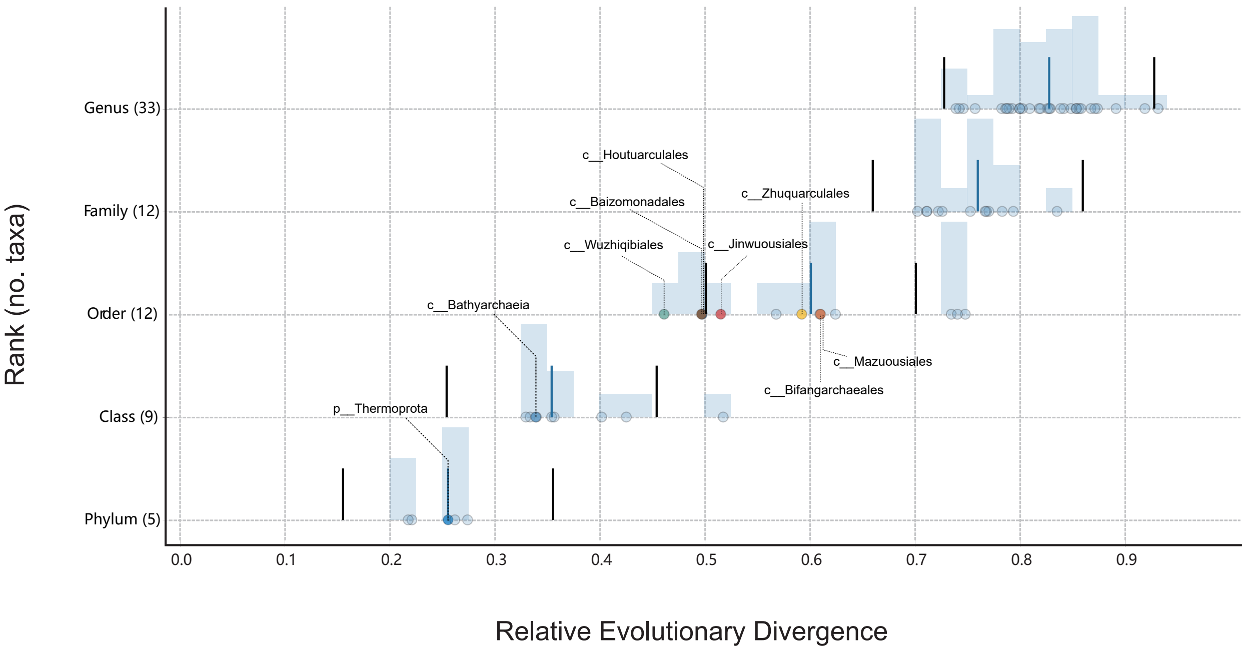


**Fig.S1.**

**The calculated relative evolutionary divergence (RED) value at each newly-proposed taxonomic rank from phylum to genus.** RED intervals for normalizing taxa at taxonomic ranks was operationally defined as the median RED value (indicated by a blue bar) at each rank ±0.1 (indicated by black bars) based on the threshold used in GTDB Release 06-RS202. Eight newly-proposed order-level units of class Bathyarchaeia are marked in different colors.

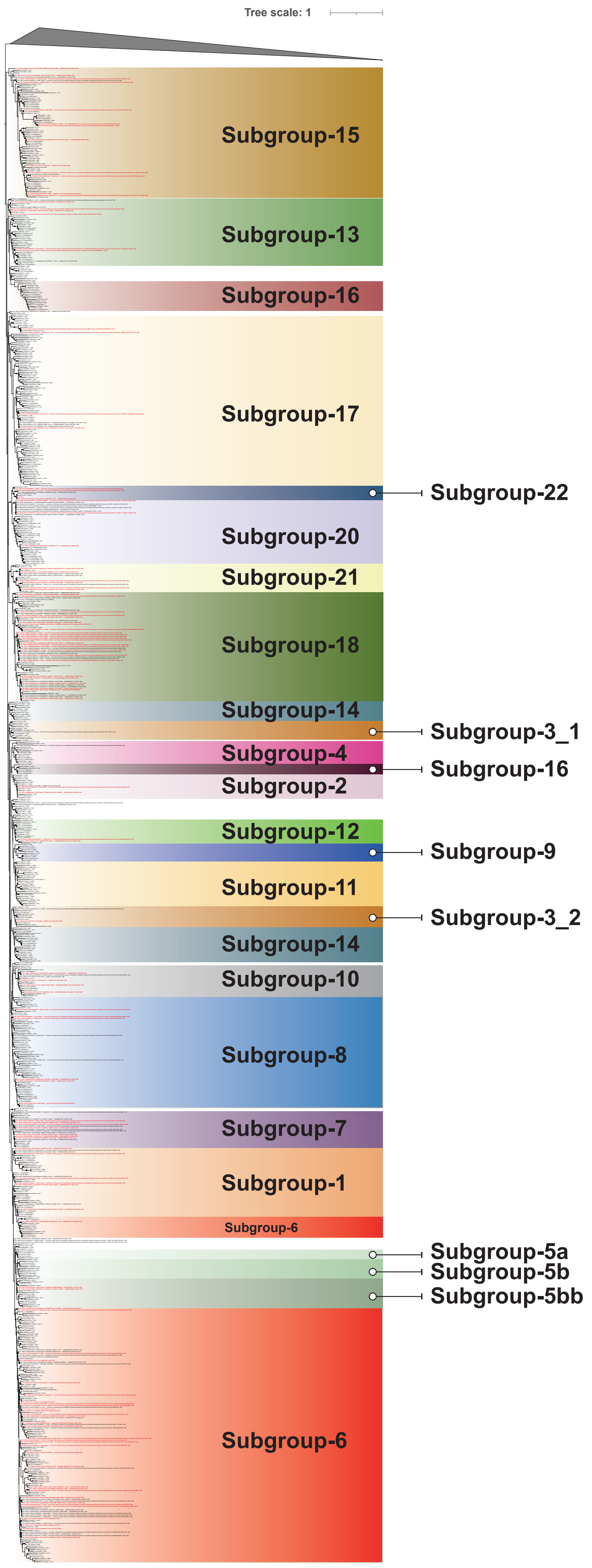


**Fig.S2.**

**The phylogenetic tree of Bathyarchaeia 16S rRNA gene.**

The maximum likelihood phylogenetic tree was constructed by using raxML version 8.2.12 with -m GTRGAMMA -N autoMRE parameter. Those red sequences indicate the 16S rRNA genes retrieved from the representative Bathyarchaeia MAGs analyzed in this study. The reference sequences with subgroup information are from Zhou *et al.,* 2019. The nodes with Bootstrap value > 75 are represented with black dot.

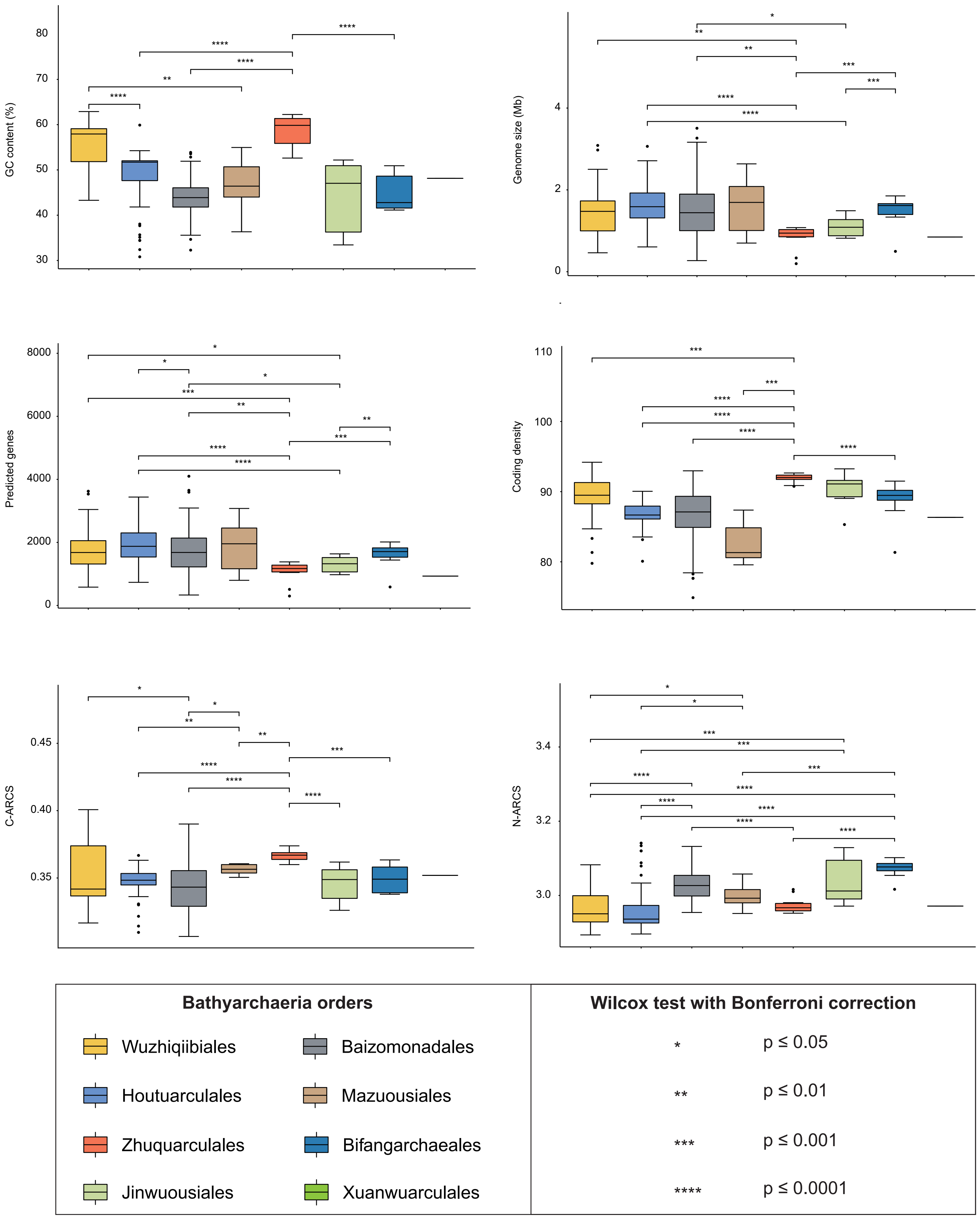


**Fig.S3.**

**Comparative analysis of the GC content, genome size, predicted gene number, coding density, C-ARCS and N-ARCS (carbon and nitrogens per amino-acid residue side chain) among the MAGs from eight Bathyarchaeia orders.**

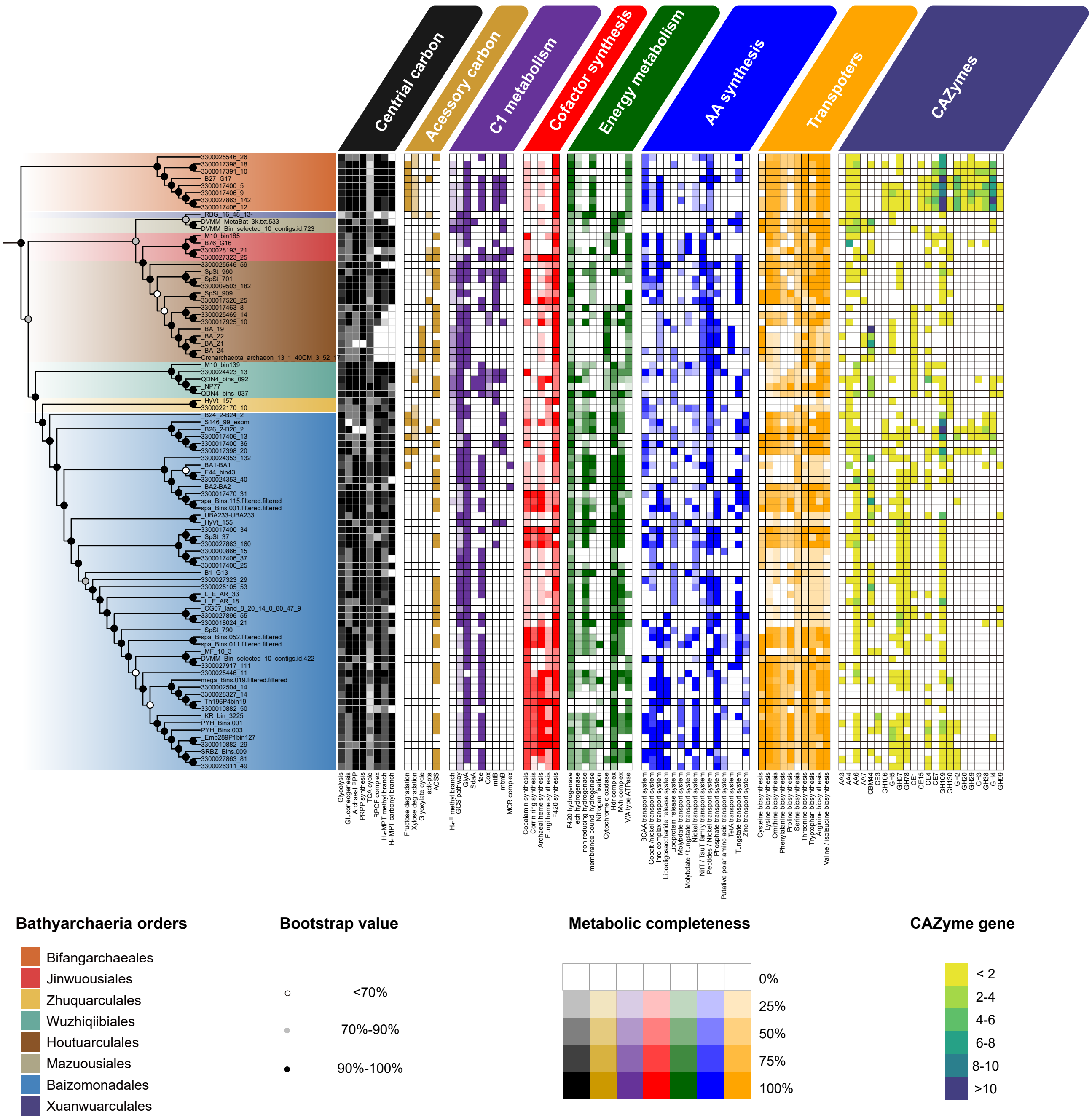


**Fig.S4 The overall metabolic potentials of 86 high quality Bathyarchaeia MAGs.**

The phylogenomic tree includes 86 high-quality representative Bathyarchaeia MAGs and was inferred by iQ-Tree2 with best-fit LG+F+R7+C20 model from 122 archaeal marker proteins implemented in GTDB-tk (see Method and Materials). The completeness of each metabolic pathway is defined by the ratio of marker genes identified in the complete gene repertoire for each MAG, while for many key genes or complexes, like those involved in C1 compounds metabolism, their presence and absence in each MAG are directly indicated by 100% and 0% completeness, respectively. Metabolic capabilities of carbohydrate degradation MAG is evaluated by the gene numbers of different CAZymes identified in each MAG. The following metabolic pathways and marker genes were used in the figure (see Supplementary Table 6): archaeal PPP, archaeal pentose phosphate pathway; PRPP synthesis, phosphoribosyl pyrophosphate synthesis; TCA cycle, tricarboxylic acid cycle; PFOR complex, pyruvate:ferredoxin oxidoreductase complex; ack-pta, acetate kinase and phosphate acetyltransferase; ACSS, acetyl-CoA synthetase; rGlyP, reductive glycine pathway; GCS, glycine cleavage system; GlyA, glycine hydroxymethyltransferase; sdaA, L-serine dehydratase; fae, formaldehyde-activating enzyme; cox, aerobic carbon-monoxide dehydrogenase; mttB, trimethylamine-corrinoid protein Co-methyltransferase; mtmB, methylamine-corrinoid protein Co-methyltransferase; MCR complex, methyl-coenzyme M reductase complex; AA, auxiliary activity; GH, glycoside hydrolase; CE, carbohydrate esterase.

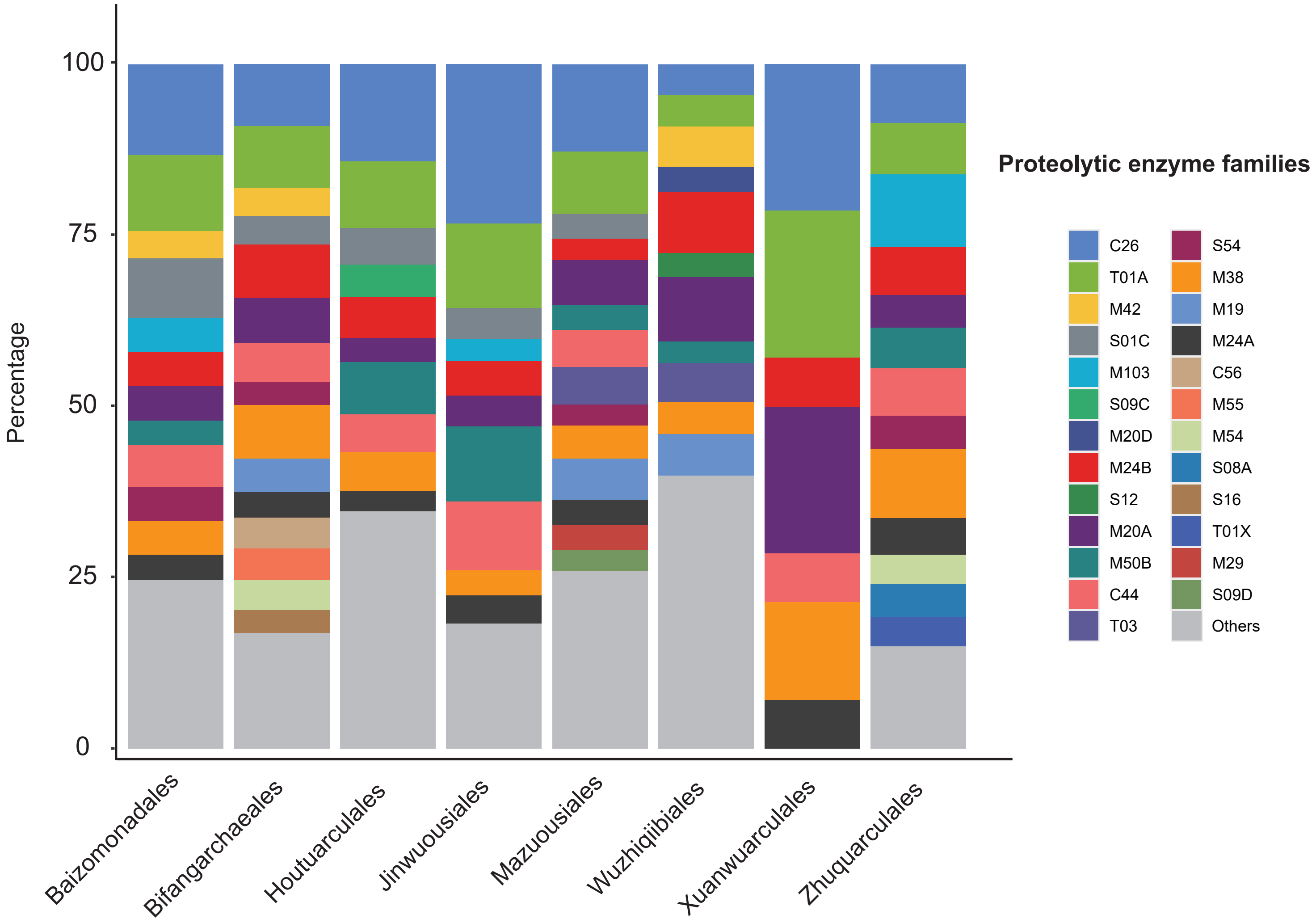


**Fig.S5.**

**The composition of different proteolytic enzyme families (based on MEROPS database) of MAGs from eight Bathyarchaeia orders.**

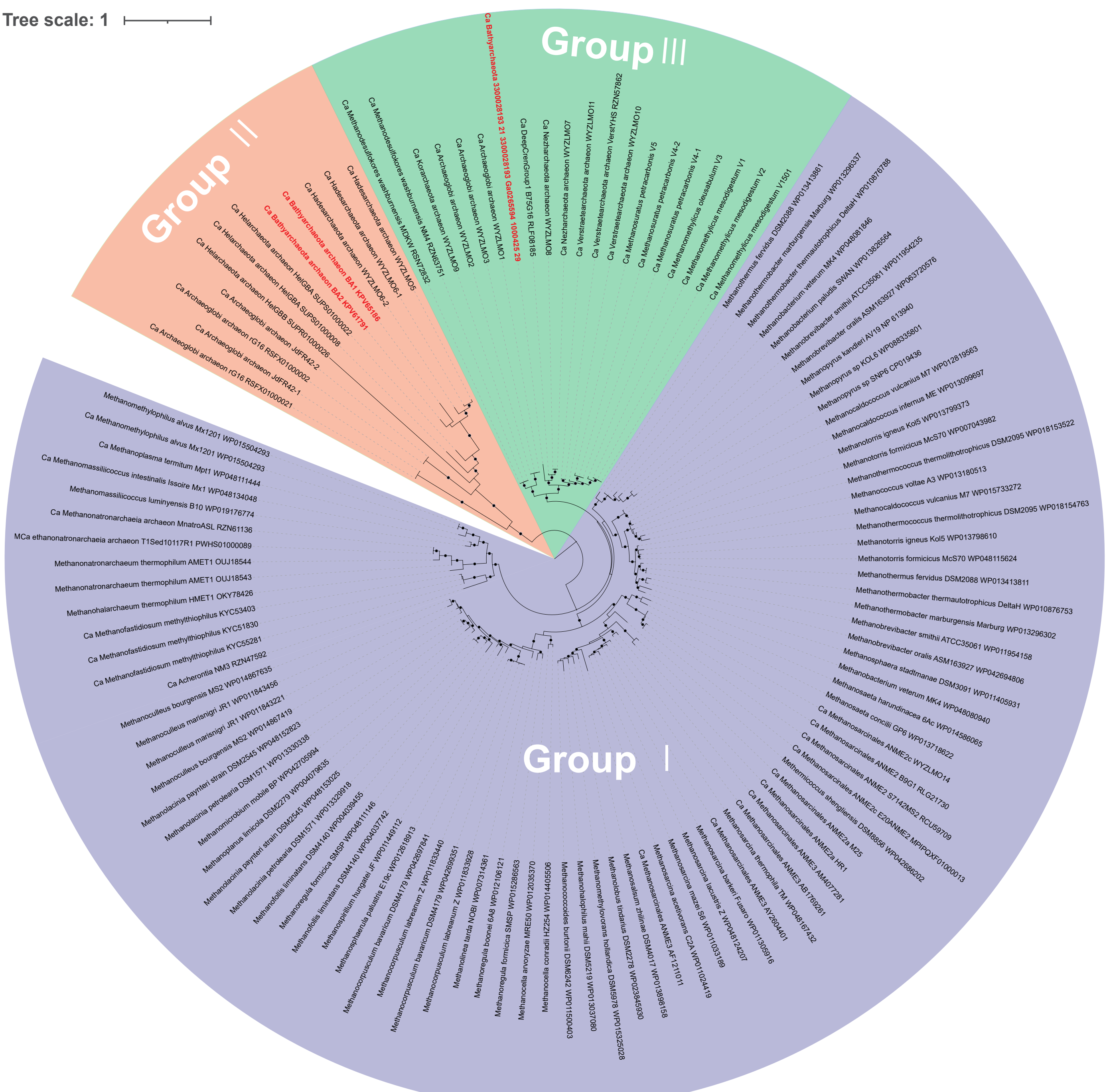


**Fig.S6.**

**Phylogenetic tree of mcrA proteins from Bathyarchaeia.**

The phylogenetic tree is constructed by IQ-TREE2 with the best-fit LG+C60+F+G model. Three mcrA-like amino acid sequences retrieved from the representative Bathyarchaeia MAGs analyzed in this study are marked in red. The reference sequences and group classification information are from Wang *et al.,* 2019.

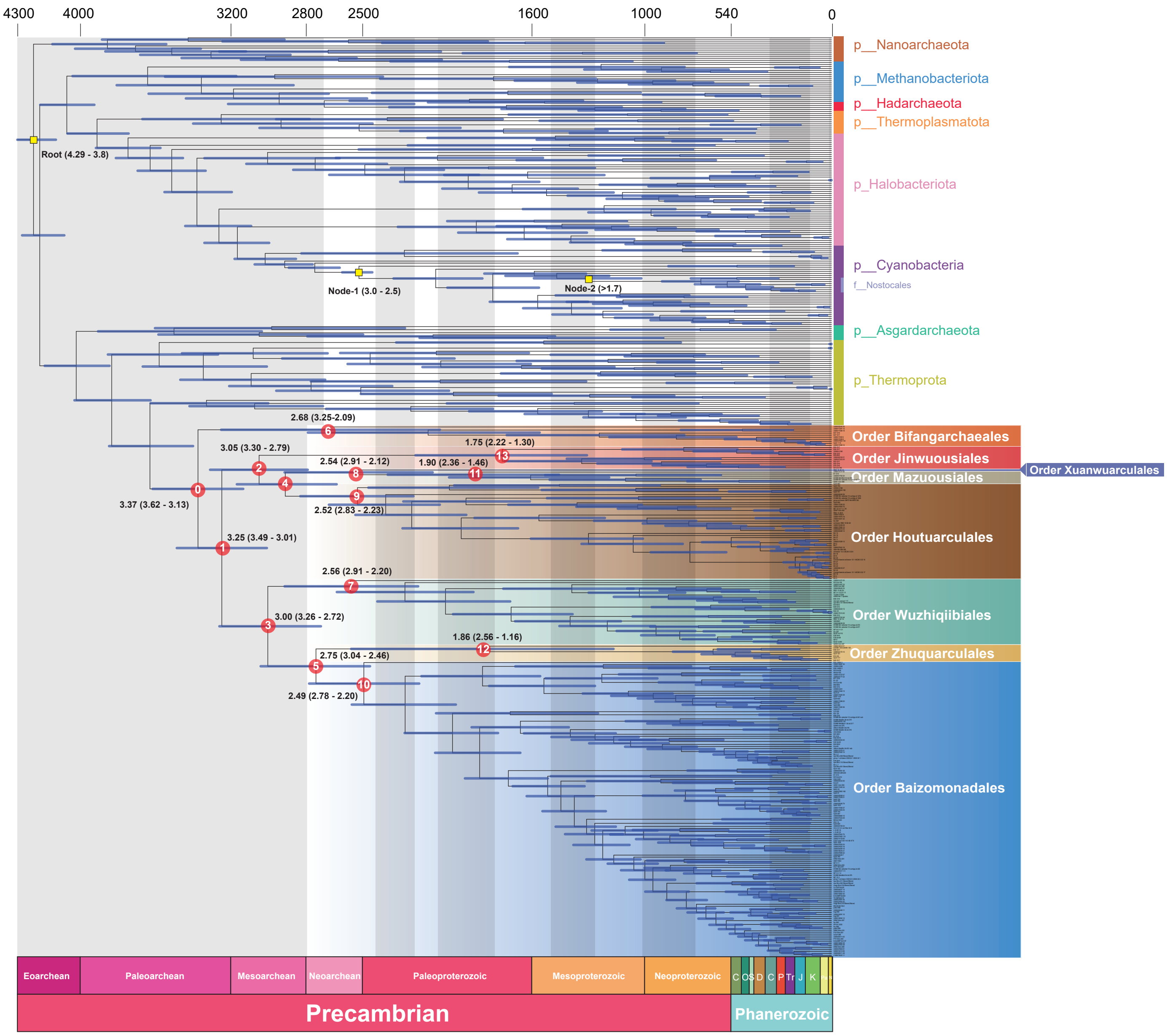


**Fig.S7.**

**Evolutionary history of class Bathyarchaeia and timing correlations with major geological activities.**

The phylogenomic tree and estimated divergence times of the major Bathyarchaeia lineages.

The whole tree was constructed based on 259 Bathyarchaeia and 190 reference MAGs by the concatenated alignment of their SMC and 16 conserved proteins using best-fit model LG+C60+F+R10 in IQ-Tree2. Molecular dating was calculated by MCMCtree with three different calibration time points (yellow squares), the timeframe displayed here is based on the calibration priors > 1.7 Ga for the lineage of Nostocales, 2.5 - 3.0 Ga for the crown group of the oxygenic Cyanobacteria and estimated divergence time of Archaea and Bacteria at 3.8 - 4.29 Ga. The divergence ages of major nodes are numbered in 1 - 11 and labeled with the posterior 95% confidence intervals (flanking horizontal blue bar).

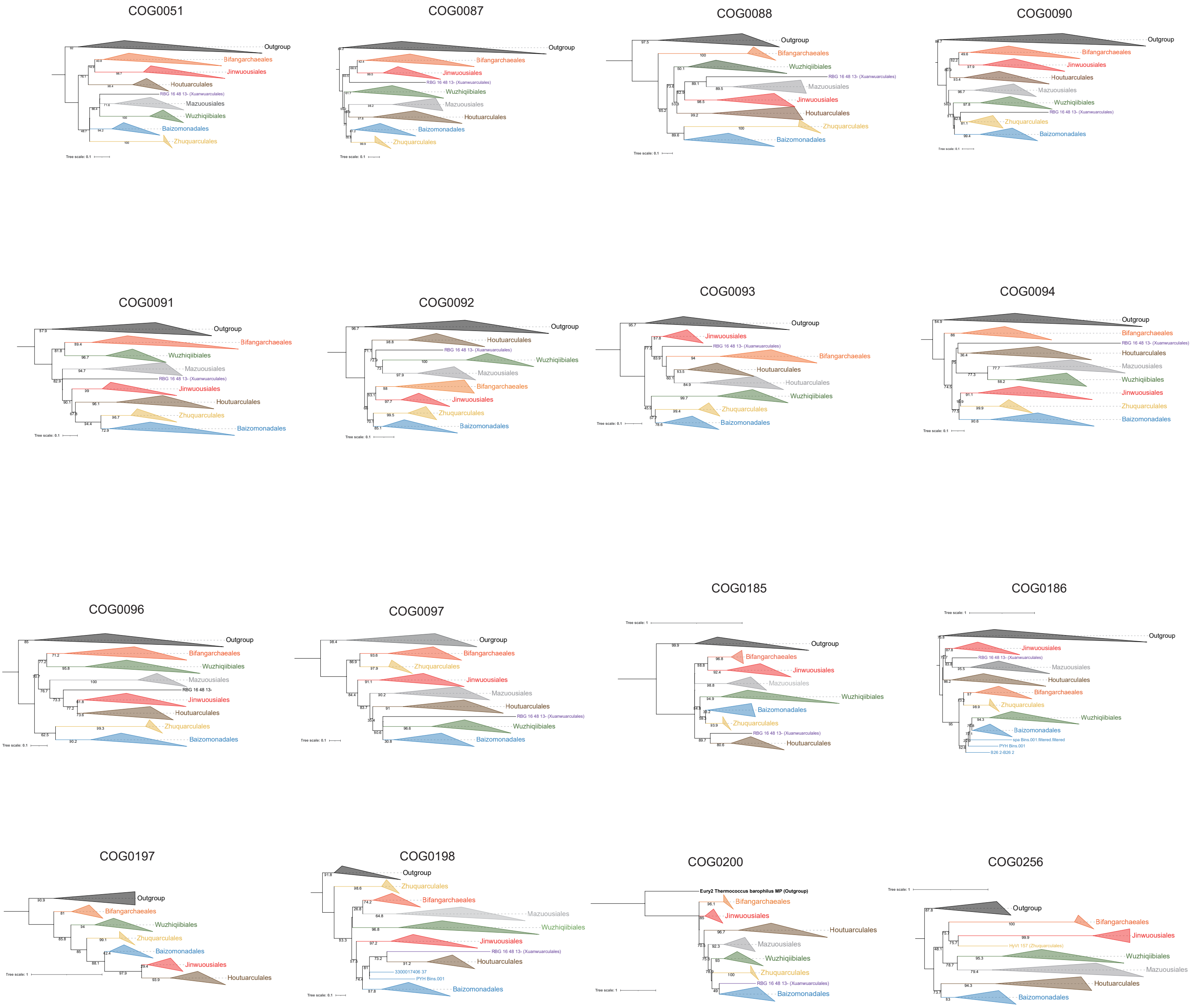


**Fig.S8.**

**The phylogenetic trees of 16 single copy conserved gene-encoding proteins used in molecular dating of Bathyarchaeia.** Each tree is build by IQ-Tree2 with their best-fit models (See Table S11)
